# Membrane Interfacial Potential Governs Surface Condensation and Fibrillation of α-Synuclein in Neurons

**DOI:** 10.1101/2025.05.30.657012

**Authors:** Jafarulla Shaikh, Anirudha N, Tuhina Mitra, Aninda Sundar Modak, Krittika Biswas, Geetanjali Meher, Aher Jayesh Bhausaheb, Nimal Archish Kannan, Bhavani Shankar Sahu, Swagata Ghatak, Sandeep Choubey, Mohammed Saleem

**Affiliations:** School of Biological Sciences, National Institute of Science Education & Research, Bhubaneshwar, India; Homi Bhabha National Institute, Mumbai, India; Center for Interdisciplinary Sciences, National Institute of Science Education & Research, Bhubaneshwar, India; National Brain Research Centre, Manesar, India; Institute of Mathematical Sciences, Chennai, India

## Abstract

Biomolecular condensates formed via liquid-liquid phase separation (LLPS) are essential for cellular organization. α-Synuclein, an amyloidogenic protein linked to Parkinson’s Disease (PD), undergoes phase separation at high concentrations, but the influence of lipid membranes on this process remains unclear. Here, combining *in vitro* reconstitution, cell biology, and simulations, we show that membranous interfaces promote α-Synuclein condensation at physiologically relevant sub-critical concentrations (∼10 nM) without crowding agents. Notably, condensation occurs only on membranes with a specific stoichiometry of lipids, underscoring the role of interfacial potential. These condensates serve as nucleation sites for fibril formation, leading to membrane deformation and rupture. A lattice gas model reveals this behavior as a prewetting-like transition, where an attractive membrane induces local phase separation below the bulk saturation concentration. Indeed altering interfacial potential by lipid composition and membrane depolarization not only drastically changes α-Synuclein puncta size and number but also triggers their release from neurons. These findings reveal the crucial role of lipid membrane interfaces in regulating α-Synuclein condensation, aggregation and release, shedding light on a potential mechanism of their cell-to-cell propagation during neurodegeneration.

## Introduction

Biomolecular condensates, also known as membraneless organelles, play a critical role in diverse cellular functions ^1^. These condensates form via phase separation of proteins, nucleic acids, and other biomolecules through weak, multivalent interactions ^2,3^. When the bulk concentration of such biomolecules exceeds a threshold—known as the saturation concentration—phase-separated condensates emerge ^4^. Classic examples include nucleoli ^5^, Cajal bodies ^6^, and stress granules ^7^. A unifying feature of many phase-separating proteins is the presence of intrinsically disordered regions (IDRs) and low-complexity domains (LCDs) ^8,9^—a characteristic also found in amyloidogenic proteins ^10–12^. α-Synuclein (α-Syn), an amyloidogenic protein associated with Parkinson’s disease (PD), has been shown to undergo phase separation at high concentrations (∼200 µM) and under crowding conditions with polyethylene glycol (PEG) ^13^ .

Lipid chemical properties—such as headgroup charge and degree of unsaturation— are known to influence α-Syn aggregation kinetics ^14^. Soluble α-Syn encounters several intracellular membranes, including the inner leaflet of the plasma membrane, synaptic vesicles, and organellar membranes. Recent studies reveal that membranous interfaces not only recruit but also regulate biomolecular condensates involved in immune responses ^15^, cytoskeletal assembly ^16^, presynaptic contact zones ^17^, RNP granules in fungi ^18^, and endocytosis ^19^. However, how membranous interfaces influence α-Syn condensate formation and their transition to fibrils— particularly under physiologically relevant protein concentrations—remains unknown.

In this study, we combine *in vitro* reconstitution, numerical simulation and live/fixed cell imaging of primary neurons, to explore how membrane composition and interfacial properties regulate α-Syn behavior. We show that membrane surfaces composed of phosphatidylcholine:phosphatidylserine (PC:PS) at a 6:4 molar ratio promote α-Syn condensation at nanomolar concentrations (∼10 nM), even in the absence of crowding agents. This finding underscores the critical role of membrane interfacial potential in driving condensation. These membrane-associated condensates act as nucleation sites for amyloid fibril formation. Using a lattice gas model, we demonstrate that this behavior resembles a prewetting-like transition, where membrane attraction drives α- Syn condensation on the membrane surface below the bulk saturation threshold ^20^.

During the formation of α-Syn condensates, membranes undergo increased rigidity. Eventually, as the condensates mature and reach a critical threshold, they rupture - releasing fibrils that cause pronounced membrane deformation. Perturbation of membrane surface charge by external addition of PS liposomes reduced the size of pre-induced α-Syn puncta, likely by inhibiting PS synthase activity and disrupting interfacial potential. On the contrary, neuronal membrane depolarization via depolarizing agents led to ∼ 10 - fold increase in α-Syn puncta as well as elevated extracellular release in primary neurons. Together, our findings reveal that membrane interfacial potential is a key regulator of α-Syn condensate formation and fibril nucleation. These condensates, in turn, can remodel and deform membranes, thus, promoting fibril propagation, potentially contributing to the pathogenesis of neurodegenerative diseases such as Parkinson’s.

## Results

### Membrane Composition Dictates Surface-Mediated Condensation Of α-Syn At Nanomolar Concentrations

α-Synuclein (α-Syn) exists in both soluble and membrane-bound forms. Notably, recent studies have demonstrated that the soluble form can undergo liquid-liquid phase separation (LLPS) ^13^. However, the role of membrane surfaces in modulating the phase separation behaviour of α-Syn remains poorly understood. To address this gap, we reconstituted purified α-Syn onto supported lipid bilayers that mimic the compositional heterogeneity of neuronal membranes. We first examined α-Syn binding and aggregation kinetics in the presence of membrane vesicles composed of total brain extract (TBE) as well as minimal model membranes made from di-oleoyl-phosphatidylcholine (DOPC) and di-oleoyl-phosphatidylserine (DOPS) at varying molar ratios (DOPC:DOPS = 1:0, 6:4, 4:6, 4:1). Strikingly, purified α-Syn displayed shorter nucleation kinetics in the presence of DOPS and DOPC:DOPS (6:4), where fibrillation was initiated around 25–27 hours (Fig. 1a & Supplementary Fig. 1). In contrast, TBE, DOPC, and DOPC:DOPS at 4:1 or 1:1 ratios exhibited nucleation kinetics similar to α-Syn alone, as reflected in the ThT fluorescence (Fig. 1a). Remarkably, the ThT signal in the DOPC:DOPS (6:4) condition, after rising, dropped sharply around 50 hours—behaviour not observed with any other membrane composition (Fig. 1a).

**Fig. 1.**
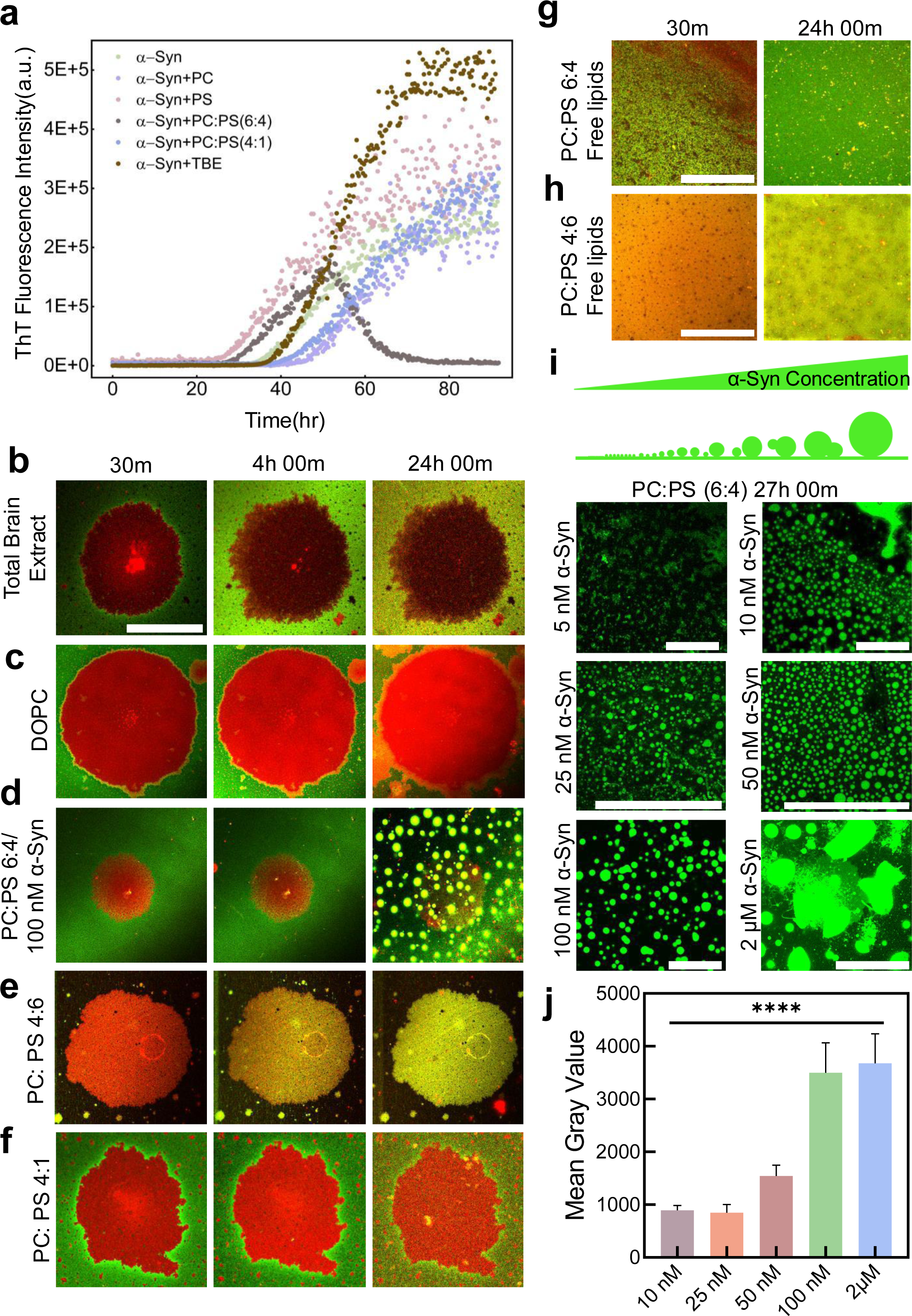
Lipid Specificity and membrane surface-induced condensation of α-Syn at nanomolar concentrations. (**a**) Thioflavin T (ThT) fluorescence assay was done to monitor the aggregation kinetics of α-Syn under shaking conditions in the presence of LUVs. α-Syn (50 μM) was incubated with 200 μM LUVs, 10 μM ThT and 0.05% sodium azide in Nap buffer, with α-Syn alone serving as control. (n=3 independent experiments). (**b-f**) SLBs of different membrane compositions labelled with Rhodamine-PE (red channel) were incubated with 100 nM α-Syn doped with Hilyte- 488 (green channel) to evaluate protein-membrane interaction over a period of 24 hours (n=3 independent replicates for each condition), Scale bar, 25 μm). (**b**) In the case of the total brain extract (TBE) membrane, only weak peripheral binding was observed. (**c**) In the DOPC membrane, there was no binding observed initially, but after 24 hours, peripheral binding to the SLB was observed. (**d**) Surprisingly, there was strong binding of α-Syn to the SLB observed in the PC:PS (6:4) membrane and at 24 hours, condensates of α-Syn were observed. (**e**) In the case where the ratio is flipped, i.e. the PC:PS (4:6) membrane, strong binding of α-Syn to the SLB, and micellar structures were observed. (**f**) No binding was detected in the case of PC:PS (4:1) membrane. (**g**) Tiny micellar structures were observed in the free-lipid condition of PC:PS (6:4) in the presence of α-Syn. (n=7 independent experiments) (Scale bar, 25 μm) (**h**) No condensate formation was observed in the case of PC:PS (4:6) in the presence of α-Syn. (n=3 independent experiments) (Scale bar, 25 μm) (**i**) Schematic illustrating the proposed mechanism of α-Syn aggregation: α-Syn binds to the membrane, leading to the formation of its liquid condensates through liquid-liquid phase separation (LLPS), which is influenced by the protein’s bulk concentration. α- Syn can be found in three different phases: dissolved in the bulk, adsorbed on the membrane, and condensed on the membrane. At low concentrations, the proteins get adsorbed on the membrane as a thin layer. When the concentration surpasses a critical threshold, condensed regions form. At even higher concentrations, liquid-liquid phase separation occurs, causing droplets to form in the bulk solution. Confocal microscopy images reveal the emergence of membrane-associated condensates 27 hours after treatment with α-Syn at the specified bulk concentration. At 5 nM concentration, no condensate formation was observed. To enhance the visibility of the condensate, the image contrast was selectively adjusted (Scale bar, 10 μm). (**j**) Fluorescence intensity measurements reveal that α-Syn condensate size directly depends on protein concentration, with higher condensate sizes forming at higher concentrations (100 nM & 2 μM) compared to smaller condensate sizes at lower protein concentrations (25 nM & 50 nM). Apart from this, after a certain point, the intensity difference between 100 nM & 2 μM was similar to the saturation point reached (means ± SD, n ≥ 150 droplets, ****P < 0.0001). The statistical significance was calculated using multiple Student t-tests.

To investigate the underlying mechanism, we performed long-duration time-lapse imaging of 100 nM Hilyte-labelled α-Syn binding to supported bilayers formed on cleaned glass coverslips. These experiments were conducted under evaporation-free conditions for up to 30 hours (Fig. 1b-h). While the physiological bulk concentration of α-Syn is estimated at ∼22 µM ^21^, it is known that nanomolar to low micromolar concentrations can suffice for surface-induced aggregation *in vitro* ^22^. No detectable binding of α-Syn was observed on TBE, DOPC, or DOPC:DOPS (4:1) membranes over the time-course of 25 hours (Fig. 1b, 1c, 1f & Supplementary Fig. 5, 6, 11), consistent with previous reports ^23^. However, α-Syn bound readily to DOPC:DOPS bilayers at both 4:6 and 6:4 ratios within a few hours (Fig. 1d, e & Supplementary Fig. 10). Intriguingly, condensation occurred only on the DOPC:DOPS (6:4) membrane, emerging at around 24 hours. Most notably, only 100 nM α-Syn was sufficient to drive condensation in the presence of this specific membrane composition (Fig. 1d). α-Syn did not undergo condensation at 100nM and 2 µM in the absence of membrane surface and crowding in line with previous reports (Supplementary Fig. 2). Membrane surface (PC:PS 6:4) also did not show any condensation over the time course of 26 hours in the absence of α-Syn (Supplementary Fig. 3). Aggregation of lipids in the membrane was ruled out in the absence of α-Syn, evident from no change in the ThT fluorescence (Supplementary Fig. 4).

To further probe this membrane-facilitated condensation, we examined the threshold α-Syn concentration required for condensation on the DOPC:DOPS (6:4) surface monitored over ∼27 hours (Fig. 1i). Surprisingly, the DOPC:DOPS (6:4) membrane appears to facilitate a surface condensation, whereby α-Syn formed condensates at concentrations as low as 10nM, an order of magnitude below its saturation concentration for bulk LLPS (∼200 µM) (Fig. 1i)^13^. No condensation was observed at 5 nM α-Syn (Fig. 1i). Conversely, α-Syn did not condense even at 2 µM in the absence of membranes (Supplementary Fig. 2), nor in the presence of free-floating DOPC:DOPS (6:4 or 4:6) lipids (Fig. 1g-h & Supplementary Fig. 12, 13). Notably, in classical liquid-liquid phase separation, the concentrations of the dilute and droplet phases remain constant even when the total concentration of the system is increased. In contrast, we observed that the intensity of the condensates increased with higher total concentrations, indicating a deviation from typical phase separation behaviour (Fig. 1i-j). These results indicate that surface condensation of α-Syn is highly sensitive to lipid composition. Such surface-mediated condensation is qualitatively distinct from liquid-liquid phase separation, driven by the interplay of protein-protein and protein-membrane interactions.

### Membrane Surface-Induced Condensation Of α-Syn Drives Transition To Fibrillar Aggregates

We next captured the dynamics of α-Syn condensate growth on the PC:PS (6:4) membrane surface. Initial membrane binding of α-Syn was observed around 4 hours; however, strong and stable binding became evident only after approximately 18 hours (Supplementary Fig. 7). Condensate formation began after ∼23 hours and progressively increased in size (Fig. 2a, Supplementary video 1). Widespread droplet growth across the entire field of view was observed, likely due to the accumulation of numerous lipid membrane patches and surface-associated debris over extended imaging durations (Fig. 2a, saturated red channel). Since no α-Syn binding was detected on DOPC-only membranes (Fig. 1c), we conclude that phosphatidylserine (PS) in the membrane plays a crucial role in promoting α-Syn binding and facilitating intermolecular interactions that lead to condensate formation. We further extracted the spatial dynamics over time and show that α-Syn initially forms a thin layer on the membrane surface, followed by the appearance of discrete condensates as demonstrated in kymographs (Fig. 2b). The surface condensation process appears to proceed through homogeneous mixing of α-Syn with PS lipids, followed by the emergence of an outer protein-rich layer (Fig. 2b, merge & Supplementary Fig. 14). Large condensates grew over time through Ostwald ripening, a process where smaller condensates dissolve and their material is transferred to larger ones due to surface tension effects (Fig. 2c, Supplementary Fig. 8 & Supplementary video 2) ^24^. Three-dimensional analysis of the condensates revealed an acute contact angle (Supplementary Fig. 15). Strikingly, around 26 hours, we observed fibrillar structures emerging from mature α-Syn droplets tubulating the membrane surface (Fig. 2d & Supplementary Fig. 9), providing the first direct visual evidence of the dynamics of fibril release from membrane-bound condensates. This strongly suggests that surface condensation may serve as a precursor to fibril formation. We conclude that α-Syn condensates formed on the membrane surface exhibit liquid-like properties, including diffusion, coalescence, and fusion – that are hallmarks of dynamic condensates. These structures likely act as intermediate states, wherein α-Syn initially forms liquid-like condensates that later transition into solid-like fibrillar aggregates. This highlights the dynamic nature of α-Syn aggregation and underscores a potential role for membrane-assisted biomolecular condensation as a transient intermediate in the amyloid formation pathway.

**Fig. 2.**
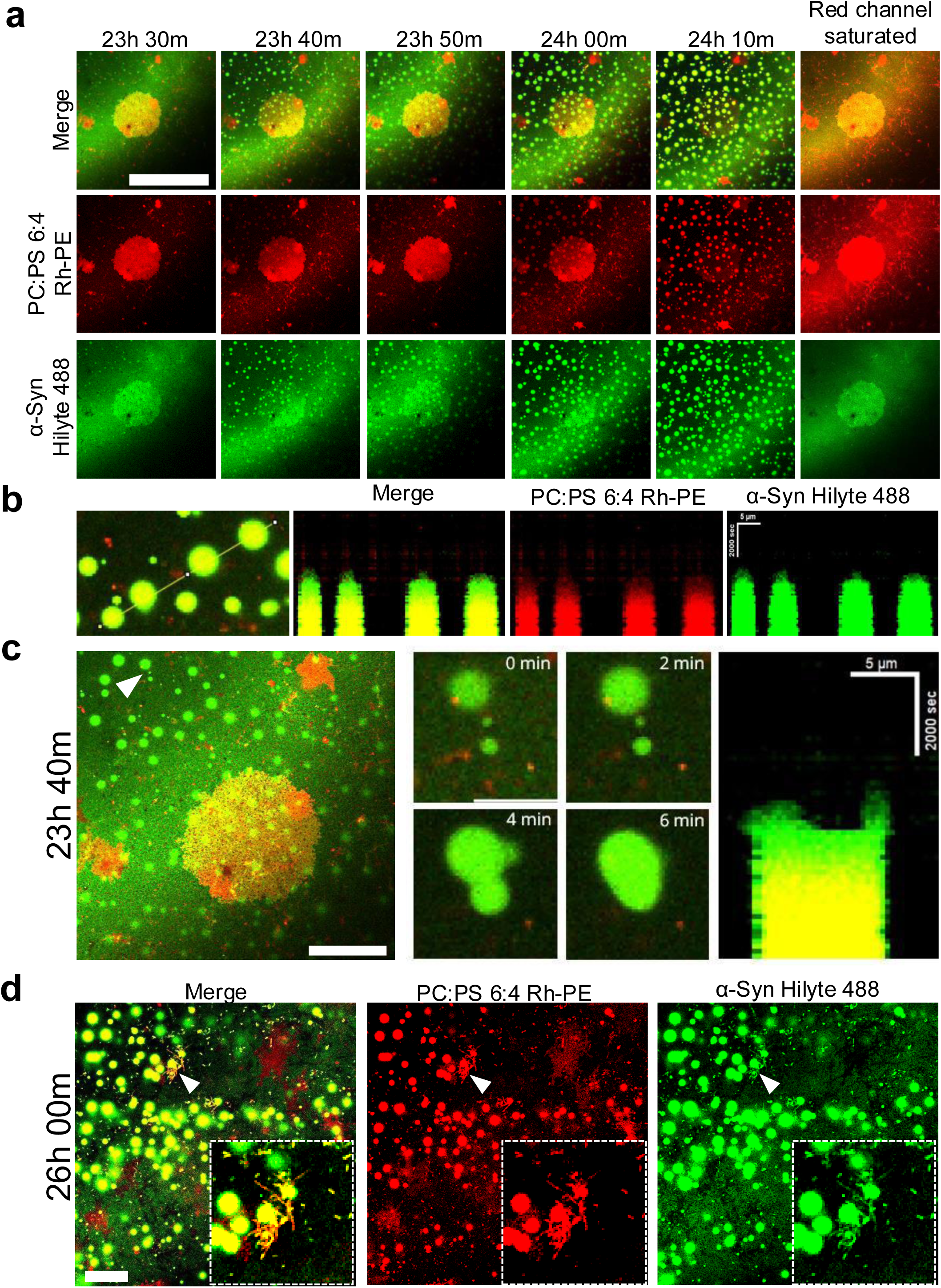
Dynamics of membrane surface induced α-Syn condensation and fibrillation. (**a**) Confocal time-lapse images show the formation of α-Syn droplets on a PC:PS (6:4) SLB. 100 nM α-Syn tagged with Hilyte-488 was incubated with Rhodamine-PE labelled SLB. (Scale bar, 25 μm.) (**b**) Kymograph reveals the temporal changes of α-Syn droplets, capturing their nucleation and progressive growth. (**c**) Time-lapse images of a fusion event of droplets after 23 hours post-incubation of α- Syn tagged Hilyte-488 with Rhodamine-PE labelled SLB. Kymograph shows the temporal progression of the fusion event. (Scale bar, 5 μm) (**d**) Confocal microscopy images show the condensates after 26 hours of incubation, and insets show fibril-like structures emanating from the droplets. (Scale bar, 20 μm.)

### Maturation of α-Syn Condensates Involves Liquid-to-Solid Transition Through Diffusion-Limited Growth

We next tracked the evolution of α-Syn condensate populations by analyzing changes in their size distribution over time. A broad range of condensate areas was observed, spanning from 6 to 65 μm². While the majority of condensates fell within the 5–30 μm² range, a smaller subset expanded to larger areas between 30–65 μm² (Fig. 3a). Further analysis revealed that the average condensate radius grew over time following a power-law scaling relationship, specifically with a time exponent of 1/3. This behaviour is characteristic of diffusion-limited Ostwald ripening (Fig. 3b). The corresponding fit also indicates that droplet volume increases linearly with time, shedding light on the dynamics and uniformity of condensate growth. To assess the internal mobility of both lipids and protein within the condensates, we performed fluorescence recovery after photobleaching (FRAP) on Rh-PE-labelled lipids and Hilyte-488-labelled α-Syn at three distinct stages of condensate maturation (i.e., 23 h 30 min, 24 h 30 min, and 25 h 30 min) (Fig. 3c-f). Mature α-Syn condensates exhibited markedly slower fluorescence recovery compared to newly formed ones (Fig. 3d), indicating a progressive reduction in mobility over time. A similar, though less pronounced, trend was observed for lipid mobility (Fig. 3f). Almost ∼65% recovery was observed for membranes without bound α-Syn indicating the fluidity of the membrane (Supplementary Fig. 16). Together, these findings suggest that α-Syn condensates undergo a liquid-to-solid transition during maturation. This gradual loss of mobility and increasing rigidity likely represents a key intermediate state in α-Syn aggregation, potentially linking phase separation to the formation of pathogenic fibrillar assemblies.

**Fig. 3.**
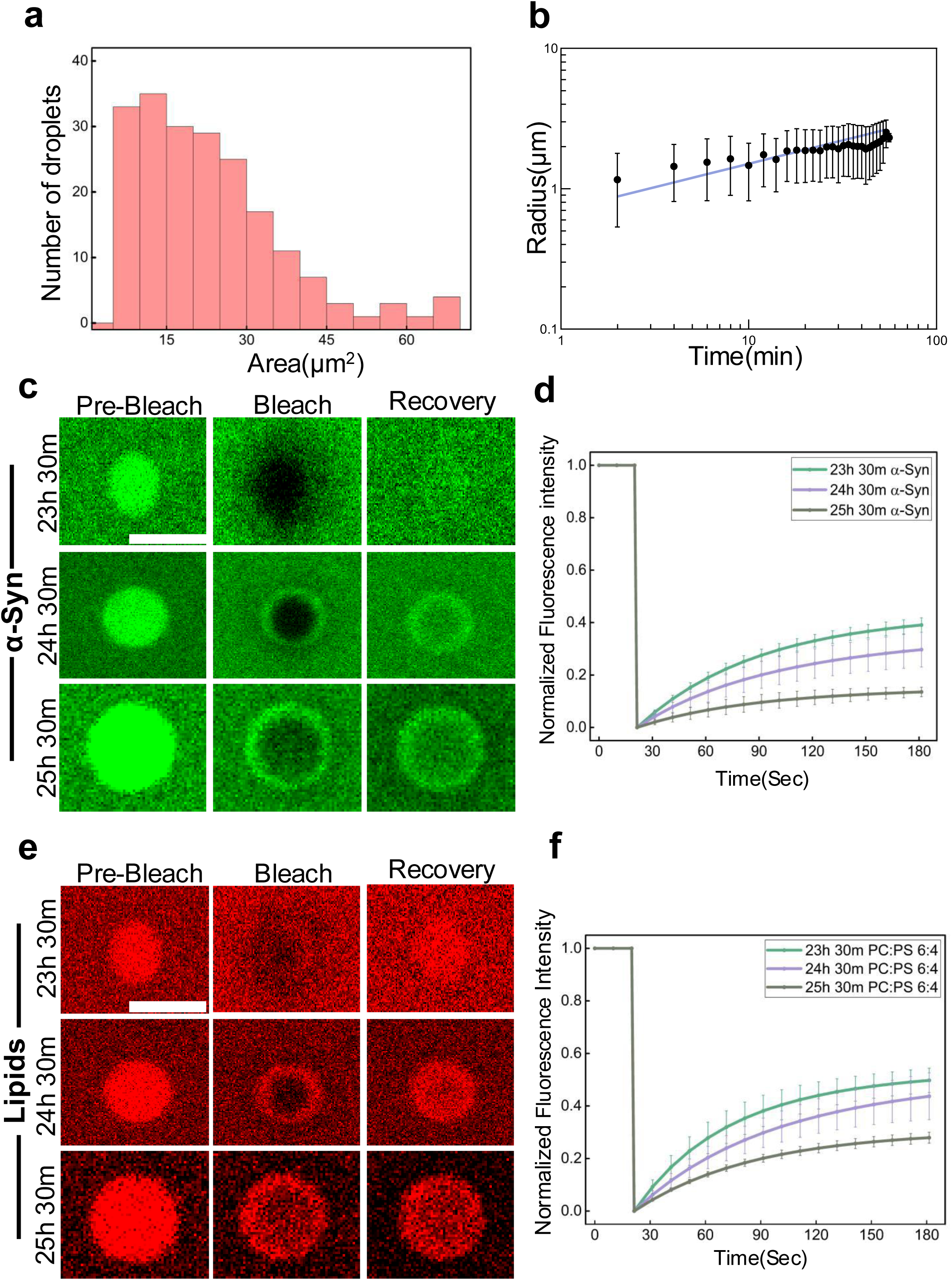
α-Syn condensates undergo liquid-like to solid-like transition. (**a**) The histogram shows the area and number of droplets at 24.40 hr timepoints in the presence of 100 nm α-Syn. (**b**) Temporal evolution of Droplet Radius (means ± SD, n=40). (**c**) Confocal microscopy images capture α-Syn aggregates during the FRAP experiment in the α-Syn imaging channel (Scale bar, 5 μm). (**d**) FRAP recovery dynamics were analysed to determine their dependence on the maturation stage of α- Syn condensates, as monitored in the α-Syn imaging channel. (means ± SD, n=3 independent experiments) (**e**) Confocal microscopy images also depict the ageing of α-Syn aggregates during the FRAP experiment in the lipid imaging channel (Scale bar, 5 μm). (**f**) The FRAP recovery behaviour was also evaluated in relation to the maturation stage of α-Syn condensates, as observed in the lipid imaging channel. (means ± SD, n=3 independent experiments).

### α-Syn Binding Reduces Membrane Tension And Compressibility

We next investigated how the membrane is affected during the stages of α-Syn binding and subsequent fibrillar growth. To probe early molecular interactions (within 1 hour), we employed Langmuir monolayer studies using surface pressure–area (π–A) isotherms to assess the thermodynamic and physicochemical changes in model membranes upon α-Syn interaction. The π–A isotherms for PC:PS (6:4) membranes exhibited a rightward shift upon α-Syn incorporation, indicating an increase in molecular area and suggesting lipid expulsion from the monolayer (Fig. 4a). A distinct plateau between 28–32 mN/m was observed, corresponding to a transition from liquid-expanded (LE) to liquid-condensed (LC) states. To quantify membrane elasticity, we calculated the compressibility modulus (Cs⁻¹), which reflects the in-plane rigidity of the monolayer. In the surface pressure range of 25–30 mN/m—relevant to bilayer mechanics—α-Syn binding caused Cs⁻¹ to rise from ∼90 mN/m to ∼180 mN/m, indicating increased membrane elasticity and reduced compressibility as the membrane entered a more ordered LC phase (Fig. 4b).

**Fig. 4.**
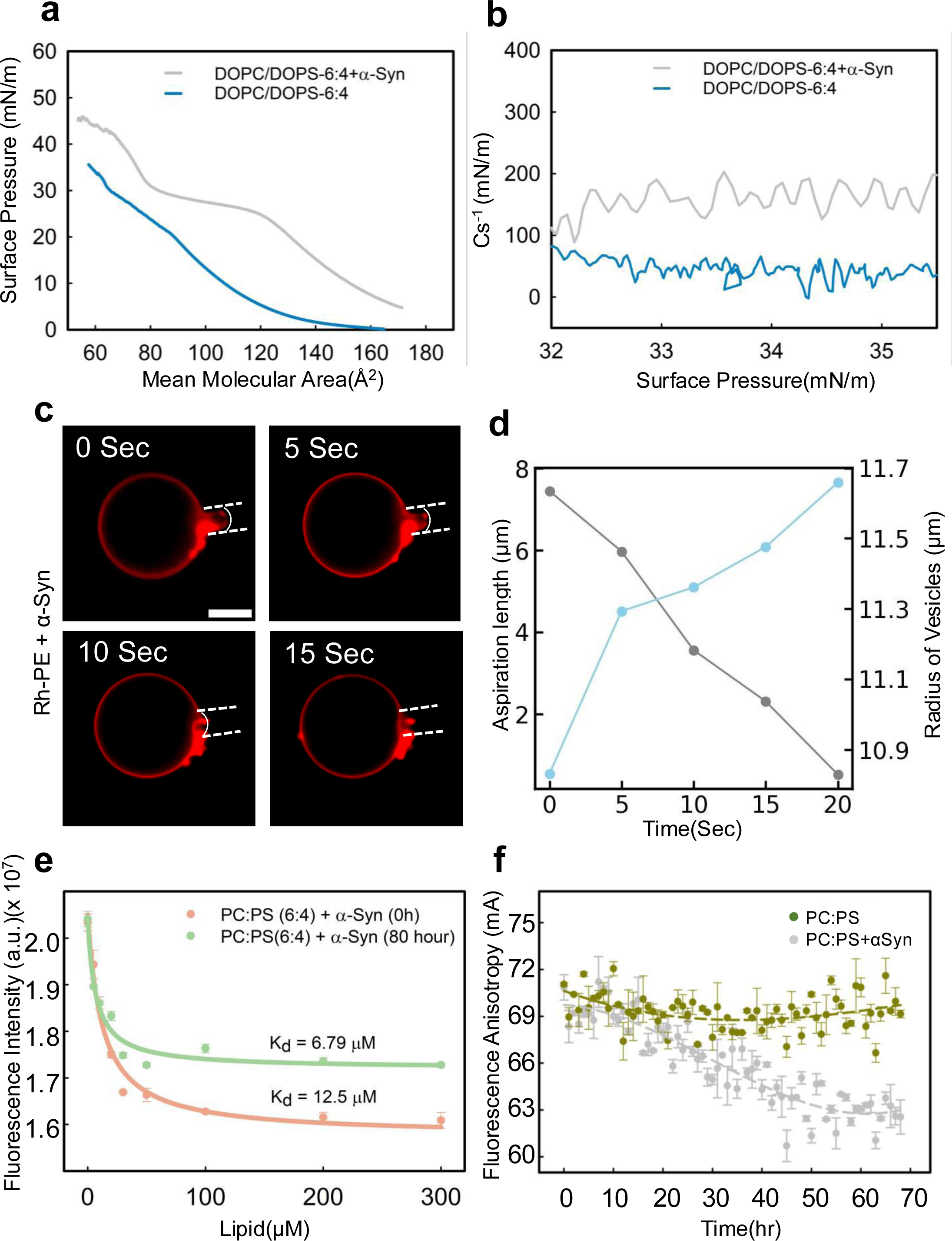
The binding of α-Syn decreases the membrane tension and compressibility modulus. (**a**) Surface pressure (π)-mean molecular area isotherm of the PC:PS (6:4) membrane was measured in the absence (blue) and presence (grey) of α-Syn. (**b**) The compressibility modulus (C_s_^-1^)-surface pressure(π) curves for the control membrane model and in the presence of α-Syn are presented in the graph. The data follows the same labelling order and colour scheme as stated in (**a**). All monolayer experiments were conducted on a PBS subphase (pH 7.4) at 25°C, with each isotherm representing the mean of three independent replicates. (For **e**, **f** and **h**, the statistical significance was calculated using one-way analysis of variance (ANOVA) followed by a Student–Newman–Keuls post hoc test with a 95% confidence interval). (**c**) Representative time-lapse Epi-fluorescence images of a GUV mimicking PC:PS (6:4) composition labelled with 0.1% Rhodamine-PE (red). Aspirated GUVs with stable protrusion length were observed. Upon injection of protein, change in spherical geometry and protrusion length was monitored over time. (n = 3 independent experiments) (Scale bar, 10μm). (**d**) Plot for time vs protrusion length (ΔLP) and the vesicle radius (R_v_). Data points are shown for a GUV (n = 3 independent experiments). (**e**) α-Syn binding to PC:PS (6:4) membrane intensifies over time. Over 80 hours, α-Syn exhibited a gradual yet striking increase in binding to PC:PS (6:4) membranes. While initial interactions were minimal, by 80 hours, the binding had surged significantly, highlighting a time-dependent strengthening of membrane association. (**f**) The addition of α-Syn caused a significant increase in the fluorescence anisotropy of TMA-DPH-labeled PC:PS (6:4) membranes, which implies that α-Syn reduces the membrane fluidity.

To examine membrane mechanics, we used micropipette aspiration of giant unilamellar vesicles (GUVs). In the absence of protein, the protrusion length (L_p_) of aspirated GUVs remained stable for over 10 minutes, establishing baseline tension (Fig. 4c, Supplementary video 3). However, after injection of 100 nM monomeric α- Syn, a reduction in L_p_ and expansion of the spherical region of the GUVs were observed, indicating a rapid drop in membrane tension. Within ∼15 seconds, GUVs relaxed and exited the pipette, undergoing notable morphological rearrangements (Fig. 4c-d, Supplementary video 4). We then explored whether α-Syn’s membrane affinity changes over the course of aggregation. Steady-state fluorescence spectroscopy revealed a significantly stronger binding at later fibrillar stages (K_d_ = 6.79 μM) compared to early-stage binding (K_d_ = 12.5 μM) (Fig. 4e), suggesting increased membrane affinity with aggregation progression. Finally, we assessed changes in membrane order using fluorescence anisotropy with the TMA-DPH probe, which reports on interfacial lipid packing. Upon α-Syn binding, the PC:PS (6:4) membrane exhibited increased fluorescence anisotropy, indicating enhanced ordering at the membrane interface. In contrast, membranes showed a progressive decrease in anisotropy over time, consistent with increasing disorder in the absence of the protein (Fig. 4f). Together, these findings demonstrate that α-Syn binding and fibrillation actively remodel membrane structure and mechanics across its aggregation timeline.

### Interfacial Potential Drives Lipid Stoichiometry-Dependent Surface Condensation of α-Syn

The interfacial potential of the cell membrane-arising from lipid composition and ion adsorption - is a critical factor essential for both intra- and intercellular interactions. We hypothesized that the lipid stoichiometry-specific binding and condensation of α- Synuclein (α-Syn) at nanomolar concentrations is governed by this interfacial potential. To test this, we measured the zeta potential of various membrane surfaces with differing phosphatidylcholine (PC) and phosphatidylserine (PS) ratios. Zeta potential, which reflects the electric potential at the shear plane (∼1–10 nm from the surface), serves as an effective proxy for interfacial potential and is determined by surface charge and the structure of the electric double layer ^25^. Our measurements revealed that α-Syn binding occurred on membranes with zeta potentials in the range of -12 mV to -20 mV, with condensation observed only at ∼ -20 mV (Fig. 5a), suggesting a threshold interfacial potential is necessary to drive surface condensation.

**Fig. 5.**
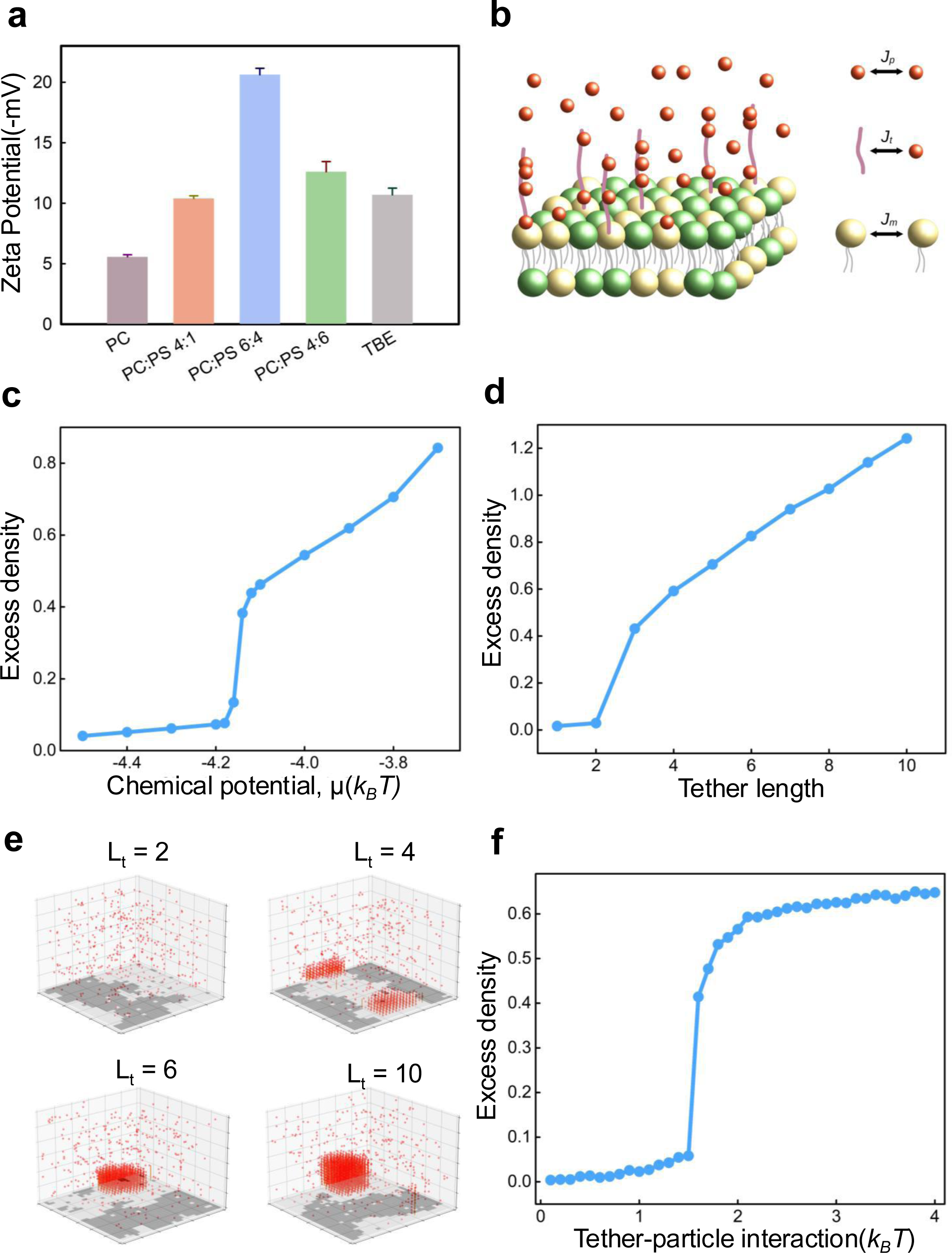
Interfacial potential drives surface condensation of α-Syn. (**a**) Changes in zeta potential were evaluated across membranes containing neutral and negatively charged compositions. (**b**) Model schematic representing the interaction of 𝛼-synuclein with the membrane. The membrane is composed of two kinds of lipids (DOPC and DOPS) and is represented via a 2D Ising model (lipid-lipid interaction J_m_). 𝛼-synuclein proteins are represented using a 3D lattice-gas model (protein-protein interaction J_p_). The proteins interact with the membrane via tethers (protein-tether interaction J_t_). (**c**) Here, we show the excess density, which quantifies the degree of surface condensation, as a function of the bulk chemical potential. We see a sudden jump in the excess density, which is characteristic of a prewetting transition. Bulk phase separation occurs at 𝜇=-3.6. (Figure parameters: J_p_=1.2, J_m_=0.5, J_t_=1.5, L_t_=5, ϕ_m_=0.5, ϕ_t_=0.2). (**d**) Excess density for various tether lengths. Surface condensation occurs only above a certain threshold tether length (L_t_=3 in this case), emphasizing the role of the range of protein-membrane interaction. (Figure parameters: J_p_=1.5, J_m_=0.5, J_t_=1.5, 𝜇=-4.6, ϕ_m_=0.5, ϕ_t_=0.2). (**e**) Simulation snapshots for various tether lengths, L_t_ = 2,4,6,10 (all other parameters same as in D). (**f**) Here, we show the excess density as a function of the protein-tether interaction strength. Similar to the tether length, there is a threshold interaction strength below which surface condensation is negligible. (Figure parameters: J_p_=1.2, J_m_=0.5, L_t_=5, 𝜇=-4.3, ϕ_m_=0.5, ϕ_t_=0.2)

To further investigate the role of interfacial potential and emergence of surface condensates, we employed a lattice-gas model, adapted from Rouches et al. ^26^ in which proteins interact with a membrane represented as a 2D Ising model tethered to a 3D lattice (Fig. 5b; see Methods). The model membrane consists of DOPC and DOPS, where α-Syn, with its positively charged N-terminal region, preferentially associates with the negatively charged PS lipids. In this framework, lipid phase separation occurs when the interaction strength between like lipids (J_m_) exceeds a critical threshold (J_cm_). Similarly, protein condensation into bulk liquid-like phases requires that the protein-protein interaction strength (J_p_) exceeds a critical value (J_cp_), provided the protein concentration surpasses the saturation limit. Importantly, even when bulk phase separation is not favored, membrane interactions can induce surface condensation via a *pre-wetting transition*, characterized by a sharp increase in protein surface density indicating the formation of a thick protein layer on the membrane (Fig. 5c). The model also predicts that increasing the protein concentration or the tether length representing the spatial range of membrane interaction—enhances surface condensation (Fig. 5c-d). Notably, below a critical tether length, surface condensation does not occur (Fig. 5e), emphasizing the role of membrane proximity and coupling strength. Additionally, a threshold interaction strength between the protein and the membrane tether is required; below this value, condensation is absent (Fig. 5f). Together, these findings underscore that membrane interfacial potential is a key driver of α-Syn condensation at concentrations below bulk saturation concentrations.

### Perturbing Membrane Interfacial Potential Via Altering Phosphatidylserine Turnover In Hippocampal Neuronal Cells Decreases α-Syn Condensate Size

Membrane interfacial potential can be modulated by altering the turnover of charged lipids at the cell surface ^27^. To test whether such perturbations influence the morphology of pre-formed α-Syn (α-Syn) aggregates or puncta, we investigated the effect of modulating phosphatidylserine (PS) levels on α-Syn aggregation in doxycycline-inducible HT22 mouse hippocampal neuronal cells.

Cells were treated with liposomes composed of PS at increasing concentrations (70 μM, 100 μM, and 150 μM) for 24 hours (Fig. 6a). Immunofluorescence imaging using an anti-Myc antibody to detect α-Syn revealed a dose-dependent reduction in puncta size, with the most pronounced decrease observed at 150 μM PS (Fig. 6b). This reduction in α-Syn puncta size is likely a consequence of inhibited PS synthase activity. When cells are exposed to exogenous PS, the internal production of PS is suppressed via feedback inhibition of PS synthase—the enzyme responsible for endogenous PS biosynthesis ^28,29^. As a result, the altered lipid composition of the plasma membrane reduces the interfacial potential, which in turn modulates membrane-protein interactions and suppresses surface-associated α-Syn condensation. These findings suggest that perturbing lipid homeostasis - specifically, through external PS treatment—can significantly alter α-Syn aggregation dynamics by modulating the membrane’s biophysical landscape.

**Fig. 6.**
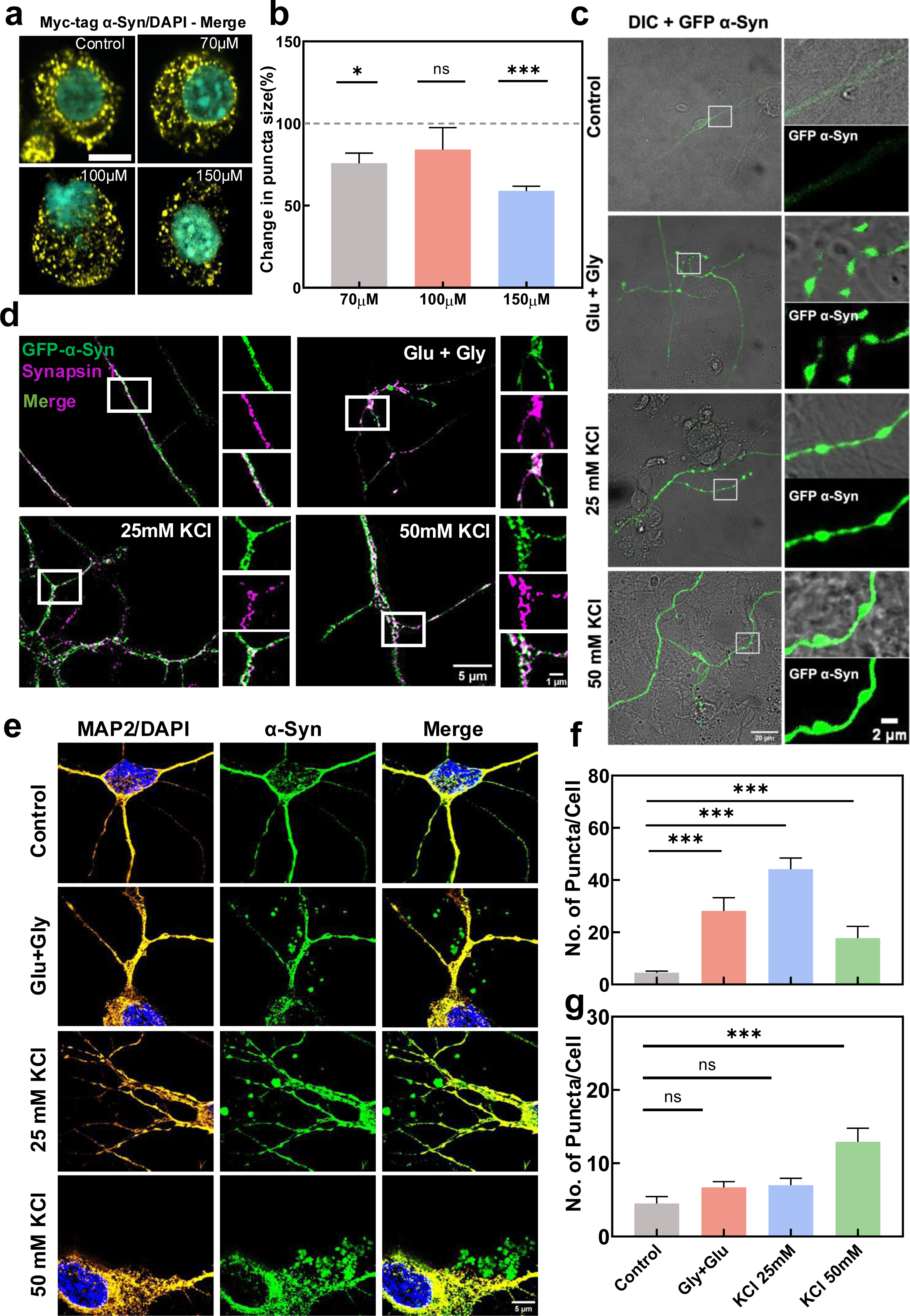
Lipid modulation and membrane depolarization alter interfacial potential, dictate α-Syn aggregation in hippocampal HT22 and neuronal cells. (**a**) HT22 cells expressing α-Syn under doxycycline induction were treated with increasing doses of Phosphatidyl Serine (PS) (70, 100 and 150 μM) for 24 hours. Immunofluorescence images, using Myc-tagged antibodies to visualise α-Syn, are shown. (Scale bar, 5 μm.) (**b**) Quantification of α-Syn aggregate size after 24 hours of PS treatment (means ± SD, n = 6 independent experiments, ns (non-significant) P > 0.05, *P < 0.05, ***P<0.001). A statistically significant reduction in aggregate size was observed at 150 μM. (**c**) Representative confocal images show α-Syn puncta distribution in neurons under resting membrane potential and after depolarisation induced by glutamate and glycine (10:1), 25mM KCl and 50mM KCl. Higher magnification insets clearly reveal an increase in puncta α-Syn upon depolarisation. (**d**) Co-immunostaining of α-Syn with presynaptic marker Synapsin 1 reveals strong co-localization in depolarized neurons, indicating recruitment of α-Syn to presynaptic compartments. Zoomed-in images further highlight the pronounced distribution of α-Syn within these presynaptic compartments upon depolarization. (**e**) Representative confocal images showing extracellular α-Syn puncta (outside neurons) at resting membrane potential and following depolarization with glutamate and glycine (10:1), 25mM KCl and 50mM KCl. An increase in extracellular α-Syn puncta is observed with increasing depolarization, suggesting enhanced release of α-Syn under stimulated conditions. (**f**) Quantification of intracellular α-Syn puncta under increasing depolarization conditions increases in depolarized neurons (glutamate and glycine (10:1), 25 mM KCl and 50mM KCl) compared to resting conditions. (**g**) Quantification of extracellular α-Syn puncta under increasing depolarization conditions reveals a positive correlation between the degree of depolarization and extracellular α-Syn puncta. Data are expressed as mean ± SEM. (n = 3 independent experiments.). For **b**, the statistical significance was calculated using a one-sample t-test with a hypothetical mean of 100, and for f and g, the statistical significance was calculated using multiple student t-tests.

**Fig. 7.**
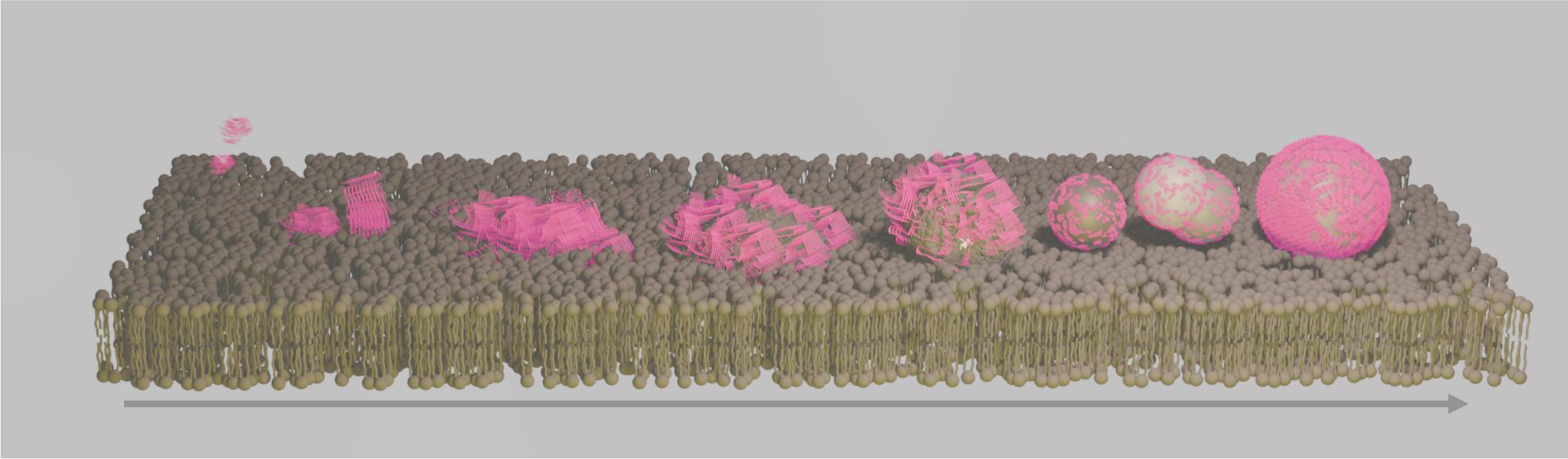
**A schematic of the proposed model illustrating membrane surface driven condensation of α-Syn and subsequent fibrillation**.

### Membrane Depolarization Drives α-Syn Condensate Formation and Extracellular Secretion in Primary Cortical Neurons

Membrane interfacial potential can also be modulated by altering ionic activity across the plasma membrane. To examine how such changes affect α-Syn (α-Syn) condensate formation, we expressed GFP-tagged α-Syn in rat primary cortical neurons and experimentally altered their membrane potential. Primary neurons are ideal for this purpose, as they undergo dynamic membrane potential changes during synaptic activity. Previous studies have linked neuronal depolarization with increased extracellular secretion of α-Syn, although the mechanisms remain unclear ^30^ ^31,32^. To investigate whether membrane depolarization modulates α-Syn interaction with neuronal membranes, we treated neurons with depolarizing agents—25 mM and 50 mM KCl, as well as glutamate. Upon depolarization, α-Syn formed punctate structures within neurons, which showed substantial colocalization with presynaptic markers (Fig. 6c-d). Quantification revealed a ∼ 10-fold increase in the number of intracellular α-Syn puncta in depolarized neurons compared to controls at resting potential (Fig. 6c, 6f). Depolarization was confirmed through calcium imaging in the same neuron–astrocyte co-cultures used for α-Syn puncta analysis (Supplementary Fig. 19a-b; Supplementary Video 5-7). The degree of colocalization between α-Syn puncta and the presynaptic marker synapsin-1 also increased in a concentration-dependent manner with KCl stimulation (Fig. 6d; Supplementary Fig. 20). Interestingly, while 25 mM KCl stimulation significantly increased intracellular puncta, no further increase was observed at 50 mM KCl. However, extracellular α-Syn puncta increased in both depolarization conditions and positively correlated with the extent of depolarization (Fig. 6e). This suggests that stronger depolarization may promote the growth and release of α-Syn condensates, possibly by enhancing their membrane interaction and subsequent secretion (Fig. 6g).

Taken together, these findings suggest that neuronal membrane depolarization— common in neurodegenerative conditions like Parkinson’s disease and dementia with Lewy bodies—can promote α-Syn condensation and facilitate its membrane association and extracellular release, potentially contributing to its pathological spread.

## Discussion

Aggregation of α-Syn is implicated in neurodegeneration; however, the dynamics of the early events that govern this process remain unclear. The lipidome of a human brain is spatiotemporally highly heterogeneous ^33^. Elevated levels of Phosphatidylserines are reported in neuronal membranes in Synucleopathies ^34^. This prompted us to investigate the role of altering lipid environment on the α-Syn aggregation. α-Syn is known to form liquid-like droplets at high critical concentrations (∼200uM) in the presence of molecular crowders and low pH ^13^. In sharp contrast, here we discover that a lipid membrane surface with a defined interfacial potential is sufficient to drive α-Syn condensation even at sub-critical concentrations (10-100nM), eventually leading to fibrillation. We demonstrate that indeed the stoichiometry of the lipid components of the membrane plays an important role in the binding of α-Syn (Fig. 1). Condensation of α-Syn is only induced in the presence of a membrane surface with specific stoichiometry of PC:PS (6:4) but not free suspended lipids of the same compositional stoichiometry (Fig.1). These observations are consistent with a prewetting-like transition driven by favourable interactions of α-Syn with negatively charged membrane surface. Upon reaching a critical concentration on the membrane surface, α-Syn undergoes a transition from a thin adsorbed layer to a thick condensed layer forming macroscopic gel-like droplets that grow over time. (Fig. 2,3). Indeed, it has been shown that changes in the conformational dynamics of disordered regions of intrinsically disordered protein such as α-Syn result in the formation of gel-like structures ^7,35,36^.

Various intracellular surfaces have been recognized for their role in promoting biomolecular condensation and potentially regulating this process ^18,37^. Originally predicted by Cahn, prewetting and wetting transition plays a crucial role in biomolecular condensation on cellular surfaces, including membranes ^20,38,39^, microtubules ^40^, actin ^41,42^, and DNA ^43^. Another recent study demonstrated that N Wasp undergoes prewetting transition to form surface condensates ^44^ on lipid membranes. One intriguing aspect of the prewetting transition is that α-Syn can undergo phase separation at sub-critical concentrations of ∼ 10 nM near a membrane surface. In contrast, bulk phase separation of α-Syn requires a much higher concentration of ∼200-500 μM, which far exceeds the physiological levels observed across different biological systems.

By reducing membrane tension and compressibility, α-Syn binding facilitates condensate formation on the membrane surface, governed by a narrow range of interfacial potential regulated by PS (Fig. 4, 5). PS is indeed one of the major functionally critical lipids in cell membranes and known to regulate membrane surface charge and protein colocalization ^27,45^. Interestingly, altering cellular PS levels by providing PS liposomes externally perturbed the size of the α-Syn puncta in the cells (Fig. 6). Spatio-temporal differences and modulation of PS levels in the membrane are likely to affect the interfacial potential that can have broad impact on transmembrane potential and membrane protein function including ion channel activity ^27^. Interestingly, neuronal hyperactivity leads to translocation of α-Syn from intracellular to the extracellular side; however, the trigger remains unknown ^31,46–48^. We propose that both lipid composition and electrical activity across neuronal membranes modulate interfacial potential, thereby promoting α-Syn binding, and subsequent fibril formation (Fig. 6). Such condensation has also been reported to spatially coordinate microtubule nucleation and branching ^49,50^. Very recently electrochemical potential of biomolecular condensates is shown to regulate their physicochemical activities ^51^. Electrochemical properties can make the condensates chemically active ^52–55^ and facilitate numerous cellular functions such as translation, stress response and cell-to-cell communication ^56–58^. Condensates that function as the nucleation centres for fibrils would facilitate deformation owing to the higher affinity for membrane surface and thus help propagation across the membrane into the extracellular side. Taken together, our results offer fundamental insights into how membrane surface and interfacial potential drive α-Syn condensation in the sub-critical nanomolar concentration range, followed by fibril-mediated membrane deformation - a process crucial to the progression of neurodegeneration.

## Materials and methods

1,2-Dioleoyl-sn-glycero-3-phosphocholine (DOPC), 1,2 dioleoyl- sn-glycero-3- phosphoethanolamine (DOPE), L-α-phosphatidylinositol (liver PI), 1,2-dioleoyl-sn-glycero-3-phospho-L-serine (DOPS), Porcine Total Brain Extract (Avanti Polar Lipids),1,2-dioleoyl-sn-glycero-3-phosphoethanolamine-N- (lissamine rhodamine B sulfonyl) (Rhod PE), and cholesterol were purchased from (Avanti Polar Lipids). Rhod PE is known to partition into liquid-disordered phases preferentially and is thus used to visualise the same ^59^. Avidin-coated chamber slides were purchased from Sigma-Aldrich.

### Protein expression and purification

α-Synuclein was expressed and purified as described earlier ^60^. Briefly, α-Syn was first expressed in E. coli using plasmid pT7-7 in BL21(DE-3)-competent cells in LB medium at 37°C until O.D. reached 0.6-0.8. they were supplemented with ampicillin to maintain plasmid selection. IPTG induction-initiated protein expression, followed by a four-hour incubation at 37°C. Cells were harvested by centrifugation and resuspended in a lysis buffer containing (50 uM Tris-HCl, 10 uM EDTA, 150 uM NaCl and PMSF, pH 7.4). Cell disruption was achieved through freeze-thaw cycles and sonication for around two hours. After boiling and centrifugation, the supernatant was treated with streptomycin sulfate (136 μL of 10% solution/mL supernatant) and glacial acetic acid (228 μL/mL supernatant) was added to remove nucleic acids. Followed by 15000 g for 10 min at four °C, Ammonium sulfate (up to 50% saturation, as calculated using the online Ammonium sulphate Calculator from Encor Biotechnology Inc.) precipitation was achieved. The precipitated protein was collected using centrifugation at 15000 g for 15 mins at 4 °C and then washed with 1 mL of 50% ammonium sulfate solution. The resulting pellet was then resuspended in 900μL of 100 mM ammonium acetate, forming a cloudy solution followed by an equal volume of ethanol at room temperature. Repetition of ethanol precipitation followed by dialysis at four °C in dialysis buffer. After the dialysis, the protein was purified using ion-exchange chromatography; the protein was loaded onto a HiTrap Q FF anion exchange column (GE Healthcare, Uppsala, Sweden). The α-Synuclein was eluted at ∼300 mM NaCl with a salt gradient from 0 mM to 1000 mM NaCl. The purification was verified by SDS-PAGE electrophoresis. Protein concentration was determined spectrophotometrically at 275 nm using a calculated extinction coefficient of 5600 M^-1^ cm^-1^.

### Thioflavin T assay for the measurement of fibrillation kinetics of α-Synuclein

To monitor the aggregation, an assay of the different α-Syn concentrations was used in phosphate buffer (pH 7.4) incubated separately with 20 μM ThT working concentration to monitor the aggregation. The lipid specificity of α-Syn was screened by incubating the protein with LUVs of different lipids for amyloid aggregation. The fluorescence measurements were performed on a Clariostar microplate reader on a Nunc black plate (Thermo Fisher Scientific). The fluorescence emission spectra were recorded on 448/482 nm excitation/emission filters. The recordings in triplicate were recorded every 15 min for 120 hours.

### Preparation of large Unilamellar vesicles

A 1 mM lipid stock was prepared and dried under nitrogen gas for each membrane condition. Then, vacuum it for an hour to remove the remaining residual solvent. The lipid film was then rehydrated with 1 mL of PBS buffer. Following a 15-minute water bath treatment, a 5-minute vortex was used. To form the large Unilamellar vesicles (LUVs), the lipid suspension was extruded using an Avanti polar extruder. Dynamic light scattering (DLS) analysis confirmed that the average size of LUV was around 130 nm.

### Preparation of Giant Unilamellar Vesicles

Giant Unilamellar Vesicles (GUVs) composed of 60% DOPC, 40% DOPS and 0.1% Rhodamine-PE were generated using the electroformation method as described in ^61,62^. To optimize GUV yield, 15 µL of a 5 mg/mL lipid mixture was uniformly spread onto indium tin oxide (ITO)-coated conductive glass slides (Nanion Technologies, GmbH) and dried under vacuum for at least 2 hours. The dried lipid film was then rehydrated in PBS buffer (pH 7.4, 300 ± 5 mOsm), and electroformation was carried out by applying a 2 V, 10 Hz sine wave for 2 hours at 60 °C.

### Preparation of supported lipid bilayers (SLBs)

The GUVs, diluted in buffer, were burst onto a plasma-cleaned coverslip at the base of a flow chamber (ibidi sticky-Slide VI 0.4) to form the supported lipid bilayers.

Following that, casein (1 mg/ml) passivation was done for 20 minutes and washed with the phosphate buffer saline before the experiment.

### Confocal fluorescence microscopy

An 8-well chamber slide from Ibidi was used for incubating the SLBs and α-Syn labelled with the Alexa-488. For GUV imaging, the slide was coated with 10 μL of 1 mg/mL streptavidin, followed by 200 μL of biotinylated GUVs, which were incubated for 30 minutes to allow immobilization. Then, the 5 μM Alexa-488 α-Syn was added to the GUV solution. The chamber was sealed with the opti-seal to prevent evaporation during the long imaging and incubation time. Imaging was conducted on a Leica TCS SP8 using the appropriate lasers for Rhodamine-PE (561nm) and Alexa-488(488nm). Identical laser power and gain settings were used during all the experiments. The image processing was done using Fiji (An image software).

### Fluorescence recovery after photobleaching

Fluorescence recovery after photobleaching (FRAP) measurements were performed on supported lipid bilayers (SLBs) doped with 0.1% rhodamine-labeled phosphatidylethanolamine(rhodamine-PE) and incubated with 100 Nm α-Syn doped with 10% Hilyte-488 α-Syn. Pre-bleach images were acquired with a low-intensity laser to establish the baseline. A specific region of interest (ROI) with a radius of 5 μm was photobleached using full laser intensity for 30 seconds, followed every few seconds for 2-3 minutes to monitor the fluorescence recovery. The experiment was repeated three times for each condition, and the resulting recovery curves were normalized to account for the variations in initial fluorescence intensity. The normalized curves were used to determine the half-recovery time (𝜏_!/#_), which is the recovery half-time. The diffusion coefficient (D_r_) was calculated using the Soumpasis equation for 2D diffusion:

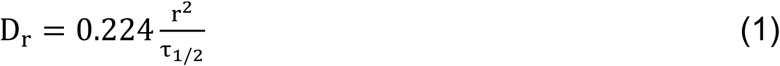

Where r is the radius of the bleached ROI (5 μm), and 0.224 is a numerically determined constant.

### Fluorescence spectrophotometric assay

The fluorescence spectroscopy experiment investigated the impact of α-Syn on lipid membrane fluidity. The Fluorescence spectroscopy experiments were conducted on previously prepared LUVs. TMA-DPH was dissolved in DMSO to reach the final concentration of 2 mM. The LUVs were then incubated with 5 μM and 500 μM TMA-DPH, bringing the total volume of the LUV mixture, α-Syn and TMA-DPH to 1 mL. Followed by 2-hour dark incubation, the fluorescence anisotropy was measured every 4 hours till 72 hours using FLS 1000 (Edinberg Instruments, UK) with excitation at 360 nm and emission at 430 nm. All the fluorescence spectra were recorded using a quartz cuvette with a fixed 1 cm path length. Anisotropy(r) was calculated automatically by the instrument using the equation:

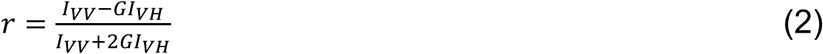

where 𝐼*_VV_* and 𝐼*_VH_*_’_ are the fluorescence intensities of the vertical and horizontal components, respectively. The grating correction factor 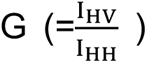 corrects for any wavelength-dependent polarizer distortion. All the experiments were repeated multiple times to ensure reproducibility and statistical significance of the results.

### Langmuir lipid monolayer experiments

Lipid monolayers were conducted using a KSV NIMA Langmuir balance equipped with two barriers for compression and a Wilhelmy microbalance with filter paper as a surface pressure sensor. The setup was enclosed in a transparent glove box to maintain a controlled environment. Before each experiment, the trough was meticulously cleaned with methanol, ethanol, and ultrapure water to eliminate surface impurities. Monolayers composed of DOPC:DOPS (6:4) were prepared by spreading a lipid/chloroform solution (1 mg/ml) dropwise, followed by a 15-mins equilibration period to allow for chloroform evaporation. The subphase consisted of phosphate-buffered saline (PBS) maintained at a constant temperature of 25°C. The monolayer was left undisturbed for 15 mins to relax the monolayer to 0 mN/m. Subsequently, a final working concentration of 100 nM α-Syn was injected into the monolayer. The solution was gently stirred using a magnetic stirrer to ensure distribution for isotherm assays. The compression was applied at a uniform 1 mm/min speed until the collapse pressure(π_c_) was reached. The isotherm data were further analyzed to calculate the compressibility modulus 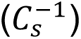, which is defined as

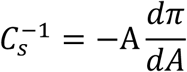

Where A is the molecular area, and 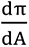 Represents the derivative of surface pressure concerning the molecular area.

### Micropipette aspiration

All the measurements were conducted using an Olympus IX83 epifluorescence microscope equipped with Hoffman modulation optics and a 100X objective lens for high-resolution imaging. Three-axis hydraulic micromanipulators were employed to position the micropipette accurately. The micropipettes were pulled from 10 μm diameter glass capillaries using a Sutter Instruments P-87 puller and a Narishige Microscope MF-900. 100 μL of GUVs was added to a custom-made open coverslip chamber. The desired suction pressure was applied to aspirate a GUV through the micropipette tip, forming an inner protrusion. Video recordings were taken over time to analyze the changes in vesicle geometry upon the addition of α-Syn.

### Generation of Stable Cell Line

Mouse hippocampal neuronal HT22 cells were transfected with 1 µg/mL of pINDUCER20-α-Syn plasmid DNA (Addgene) using Metafectene Pro (Biontex, lot no. RKP205/RK081621) following the manufacturer’s protocol. After 48 hours, the cells were transferred to 100 mm culture dishes and subjected to antibiotic selection using 7.5 mg/mL G418 (Geneticin, G-418 sulfate; GOLDBIO, G-418-25) containing DMEM medium. Selection was continued until non-transfected cells were dead, which was for 14 days in our case.

Following selection, 24 single-cell-derived colonies were isolated and seeded into 48- well plates. Upon reaching ∼80% confluency, cells were sequentially passage into 24- well, 12-well, and finally 6-well plates. Each colony was then screened for α-Syn expression via immunofluorescence using an anti-myc tag antibody, as the α-Syn construct included a myc tag. Colonies testing positive for myc-tagged α-Syn expression were selected and cryopreserved at –80 °C.

### Mammalian cell culture and treatment with monomeric α-Syn

Mouse Hippocampal Neuronal HT22 stable cells expressing α-Syn were grown in DMEM Glutamax, supplemented with 10% FBS, 1% antibiotic-antimycotic, along with 0.75mg/ml of G418 at 37° C, 5% CO2 in a humidified incubator. Early Passage cells (1-10) were used for all the experiments. Mouse HT22 doxycycline inducible stable cells were induced with 1µg/ml of Doxycycline, 24 hours after plating. Media was replaced with fresh media containing 1µg/ml of Doxycycline to which 70µM, 100µM and 150µM of exogenous Phosphatidyl Serine (DOPS), were added respectively for each condition. After 24 hours of induction and treatment with 70µM, 100µM and 150µM the cells were harvested and experiments such as Immunofluorescence Imaging were performed.

### Immunofluorescence Imaging

2*10^4^ cells were seeded in each 15 mm coverslip, and the cells were treated with exogenous Phosphatidyl Serine (DOPS) by the above-discussed protocol. After treatment, cells were gently washed with 1x PBS thrice and then fixed using 4% Paraformaldehyde (PFA) for 20 minutes at room temperature. Following fixation, cells were washed gently with 1x PBS thrice, and then the cells were permeabilized with 0.1% x Triton X-100 for 10 minutes and blocked using 1% BSA for 45 minutes. The primary antibodies, rabbit MAP2 (1:150, Cell Signaling Technology) and Guinea pig Synapsin1 (1:300, Synaptic Systems), Rabbit anti-Myc of dilution (1:1000, Invitrogen) were diluted in in the blocking solution (1% BSA) and incubated overnight at 4°C. The next day, cells were washed once with PBS and twice with PBS. Secondary antibodies, anti-rabbit Alexa Fluor 568 (1:1000, Invitrogen) and anti-Guinea pig Alexa Fluor 647 (1:1000, Invitrogen), were added in the blocking solution (1% BSA). Alexa Fluor 594 conjugated goat Anti-Rabbit (1:1000) was used, and cells were incubated for 45 minutes in a humidified chamber in the dark. Coverslips were mounted using DAPI containing mounting media, and images were captured using 100x oil objectives of Olympus IX83 Confocal Microscope.

### Primary Neuron-Astrocyte Co-culture

Cortical neurons and astrocytes were cultured from P0-P1 Sprague-Dawley rat pups. The brains were removed and placed in ice-cold calcium- and magnesium-free Hanks’ Balanced Salt Solution (HBSS) containing 10 mM glucose and HEPES (Sigma). The cortical tissue was separated and digested with 0.25% trypsin-EDTA (Gibco) and 150 units/ml DNAse (Sigma) at 37°C for 15 minutes. After digestion, trypsin activity was stopped with 10% fetal bovine serum (FBS) (Gibco). The cells were dissociated mechanically and then centrifuged at 1000 rpm for 5 minutes at 4°C. The resulting cell pellet was resuspended in culture medium composed of neurobasal-A (Gibco), 10% FBS (Gibco), 1% Glutamax (Gibco), 1% Anti-anti (Gibco), and 2% N2 supplement (Gibco). Cells were plated onto coverslips pre-coated with 0.1 mg/ml Poly-D-lysine. The cultures were maintained at 37°C in a 5% CO2 incubator with controlled humidity. The culture medium was replaced with fresh medium on day 4, and 5 µM AraC (Sigma) was added during each subsequent medium change, performed every 3 days.

### Transfection of Primary Cultures with EGFP-tagged α-Synuclein Plasmid

On day 9 in vitro (DIV9), primary neuron–astrocyte co-cultures were transfected with an EGFP-tagged α-synuclein plasmid. The plasmid DNA (0.5µg/well) and Lipofectamine™ 2000 (Invitrogen) were each diluted separately in Minimal Essential Medium (Opti-MEM™, Gibco) and incubated for 5 minutes at room temperature. The diluted DNA and Lipofectamine solutions were then combined at a 2:3 ratio (µg DNA:µL Lipofectamine) and incubated for 15 minutes at room temperature. Prior to transfection, the complete Neurobasal-A medium was removed from the cultures. The DNA-Lipofectamine complexes were added dropwise to the cells and incubated at 37°C with 5% CO₂ for 4 hours. After incubation, the medium was replaced with fresh, complete Neurobasal-A. Cells were treated after 36 hours of transfection.

### Induction of Depolarization in Transfected Primary Cells

To induce depolarization, primary cells were treated with HBSS containing 100 µM glutamate (Sigma) and 10 µM glycine (Tocris). Moreover, cells were treated with different concentrations of KCl (25 mM and 50 mM) (Fisher Scientific) in transfected cells to evaluate the exponential effects of depolarization.

### Calcium Imaging with Fluo-4 Direct™ in Primary Neuron-Astrocyte Co-culture

For the calcium assay, the loading solution was prepared by adding 10 mL of Fluo-4 Direct™ calcium assay buffer and 200 μL of 250 mM probenecid stock solution to one bottle of 2X Fluo-4 Direct™ calcium reagent (Invitrogen). The solution was mixed 1:1 with culture media to achieve a final 1X Fluo-4 Direct™ calcium reagent concentration.

After 24 hours of treatment, the co-cultures were loaded with the 1X Fluo-4 Direct™ calcium reagent. The reagent was added to the glass-bottom dish containing cells, and the cells were incubated with the solution for 30 minutes at 37°C. After incubation, the cells were incubated at room temperature for an additional 30 minutes. The dye was removed, and assay buffer was added during recording. Fluorescence intensity was recorded at 33 frames per second (fps) for 2 minutes on a Zeiss fluorescence microscope with an Axiocam 702 mono fast camera.

## Data Analysis

### Calcium Imaging Analysis

Calcium imaging data were analyzed using FIJI (ImageJ) and custom Python scripts. Regions of interest (ROIs) were manually selected for individual neurons. Fluorescence intensity (F) was recorded over time for each ROI. Baseline fluorescence (F₀) was calculated as the mean of the lowest 10 fluorescence values for each neuron. Changes in fluorescence (ΔF/F₀) were calculated using the formula: ΔF/F₀ = (F - F₀) / F₀. Peaks in calcium activity were detected using a peak-finding algorithm with parameters for minimum prominence, height, distance between peaks, and width. The data were plotted and statistically analyzed for comparisons across experimental groups.

### Colocalization Analysis

Colocalization between α-synuclein (α-syn) and Synapsin-1 was quantified using Manders’ coefficient. Images were first thresholded to isolate α-syn puncta. Particle analysis was performed in FIJI to identify and count individual α-syn puncta. Colocalization was then assessed using the Coloc2 plugin. Mander’s coefficient was calculated to measure the fraction of α-syn puncta overlapping with Synapsin-1. This Manders’ value was multiplied by the total area of α-syn puncta. This gave the absolute colocalization area of α-syn with Synapsin-1. The resulting colocalized areas were plotted for comparison across experimental groups.

### Quantification & Statistical Analysis

ImageJ software was used to analyze and process the images obtained from the experiments. A uniform detection setting was applied consistently across all the experiments to ensure comparability. The size of the condensates was manually quantified using ImageJ. Statistical analysis was performed using one-way ANOVA in SigmaPlot, with p-values less than 0.05 considered statistically significant for all the experiments. Some graphs were plotted using Origin Pro software and GraphPad (Prism).

## Acknowledgement

MS acknowledges the financial support received from the Science & Engineering Research Board (SERB) (Grant No. EMR/2017/004513), Department of Biotechnology, Govt. of India (Grant No. BT/PR/21226/MED/122/41/2016) and DBT-Wellcome Trust India Alliance Intermediate Fellowship (Grant No. - IA/I/20/2/505212). SC acknowledges Ramalingaswamy Re-entry Fellowship (Grant No. BT/HRD/35/02/2006). SG acknowledges SERB (Grant no. SRG/2022/000117), Indian Council of Medical Research (ICMR) (Grant no. IIRP-2023-0585) and DBT (Grant no. BT/PR48748/MED/122/329/2023) for funding. BSS acknowledges International Brain Research Organization, Ignite Research Foundation and DBT-NBRC for financial support. MS, SC, SG also acknowledge Department of Atomic Energy (DAE) for the intra-mural financial support. We acknowledge Vaishnav Manoj for help during Calcium imaging quantification. We also acknowledge Ashutosh Prince for his help with illustration preparation.

## Contributions

MS, SC and SG conceptualized the work; MS, SC, SG and BSS designed the research; JS, AN, TM, ASM, KB, GM, JAB performed the research; MS, SC, SG, BSS, JS, AN, TM, ASM, KB, GM, JAB, NAK analysed and interpreted the data; MS, SC, SG, JS and AN wrote the paper with contributions from all other co-authors.

## Notes

### Competing Interest Statement

The authors have declared no competing interest.

### Summary of Updates

The first author's details have been corrected on the title page of the manuscript.

