## Supplementary video legends for "Membrane Interfacial Potential Governs Surface Condensation and Fibrillation of α-Synuclein in Neurons"

### Supplementary information

**Supplementary video 1:** Dynamics of α-Synuclein (α-Syn) condensation on membrane surface (PC:PS 6:4) over a period of 24 hour 10 minutes. (Scale bar, 20 μm).

**Supplementary video 2:** Dynamics of α-Synuclein (α-Syn) droplet fusion. (Scale bar, 10 μm).

**Supplementary video 3:** Time-lapse fluorescence imaging and micropipette aspiration of Giant Unilamellar vesicles (GUVs) labeled with 0.1% Rh-PE (red). The GUVs were aspirated under constant pressure ($\Delta p$ = 50Pa) using micropipette manipulation and their morphological dynamics monitored over time. (Scale bar, 10 μm).

**Supplementary video 4:** Time-lapse fluorescence imaging and micropipette aspiration of Giant Unilamellar vesicles (GUVs) composed of PC:PS (6:4) membrane labeled with 0.1% Rh-PE (red). The GUVs were aspirated under constant pressure ($\Delta p$ = 50Pa) using micropipette manipulation in the presence of 100 nM α-Synuclein and their morphological dynamics were monitored over time. (Scale bar, 10 μm).

**Supplementary video 5:** Time-lapse calcium imaging of neuronal activity at resting membrane potential.

**Supplementary video 6:** Time-lapse calcium imaging of neuronal activity under glutamate-induced depolarization.

**Supplementary video 7:** Time-lapse calcium imaging of neuronal activity under 25mM KCl induced depolarization.
