## Supplementary dataset for "Membrane Interfacial Potential Governs Surface Condensation and Fibrillation of α-Synuclein in Neurons"

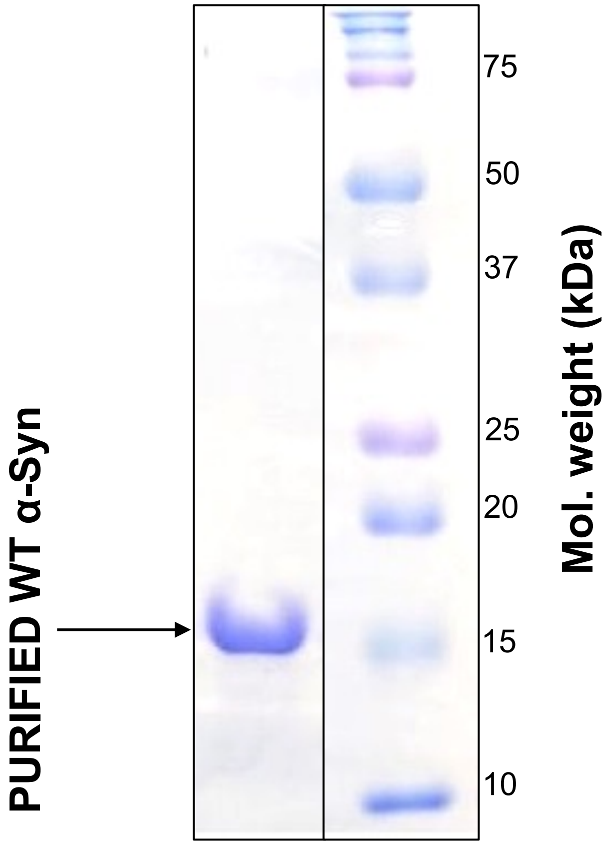


**Supplementary Figure 1: Expression and Purification of recombinant α-Synuclein (α-Syn).** SDS-Page of α-Syn, recombinantly expressed and purified from pT7-7 wild-type plasmid through anion exchange chromatography. Coomassie blue-stained gel after gel filtration shows highly pure monomeric α-Syn, with ~90% purity by densitometry measurement.

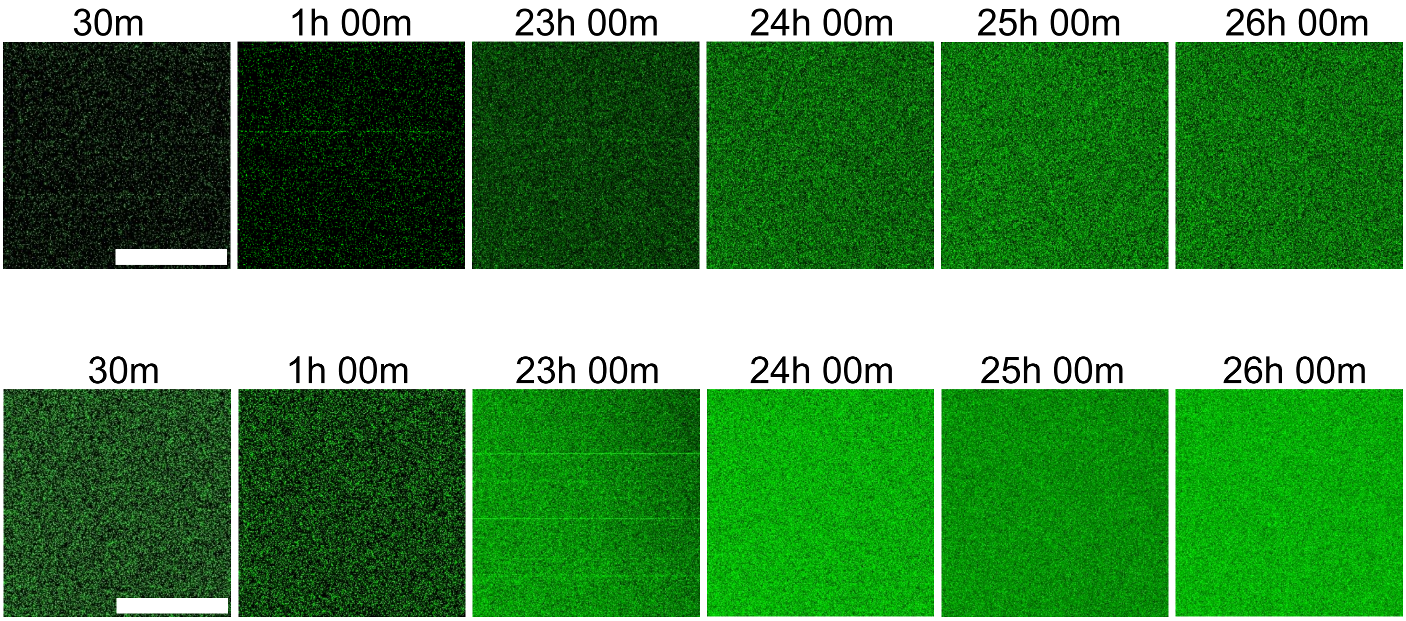


**Supplementary Figure 2: α-Syn does not undergo condensation in the absence of the membrane surface.** No condensate formation was observed in the submicromolar (100nM) and high concentration (2μM) of α-Syn in PBS buffer in the absence of lipids during confocal microscopy imaging over a 26-hour period. (Scale bar, 20 μm). n = 3 independent experiments.

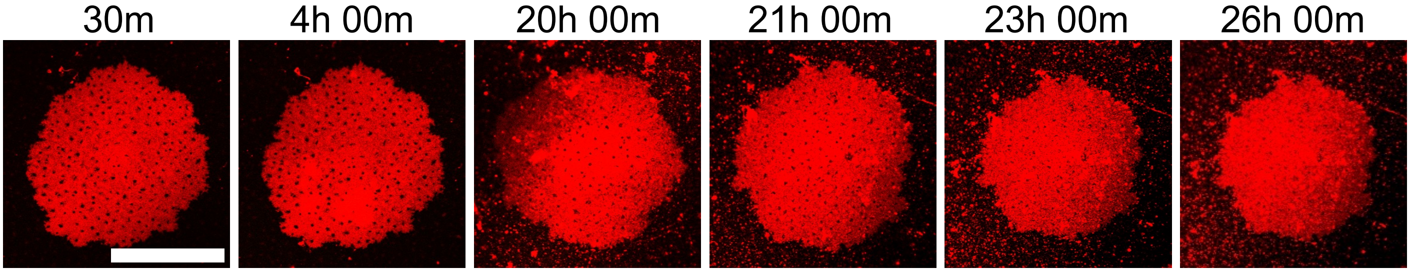


**Supplementary Figure 3: Membrane surface in the absence of α-Syn.** Confocal microscopy imaging over a 26-hour period showed no formation of condensates in the PC:PS (6:4) SLB in the absence of α-Syn. (Scale bar, 20 μm). n = 3 independent experiments**.**

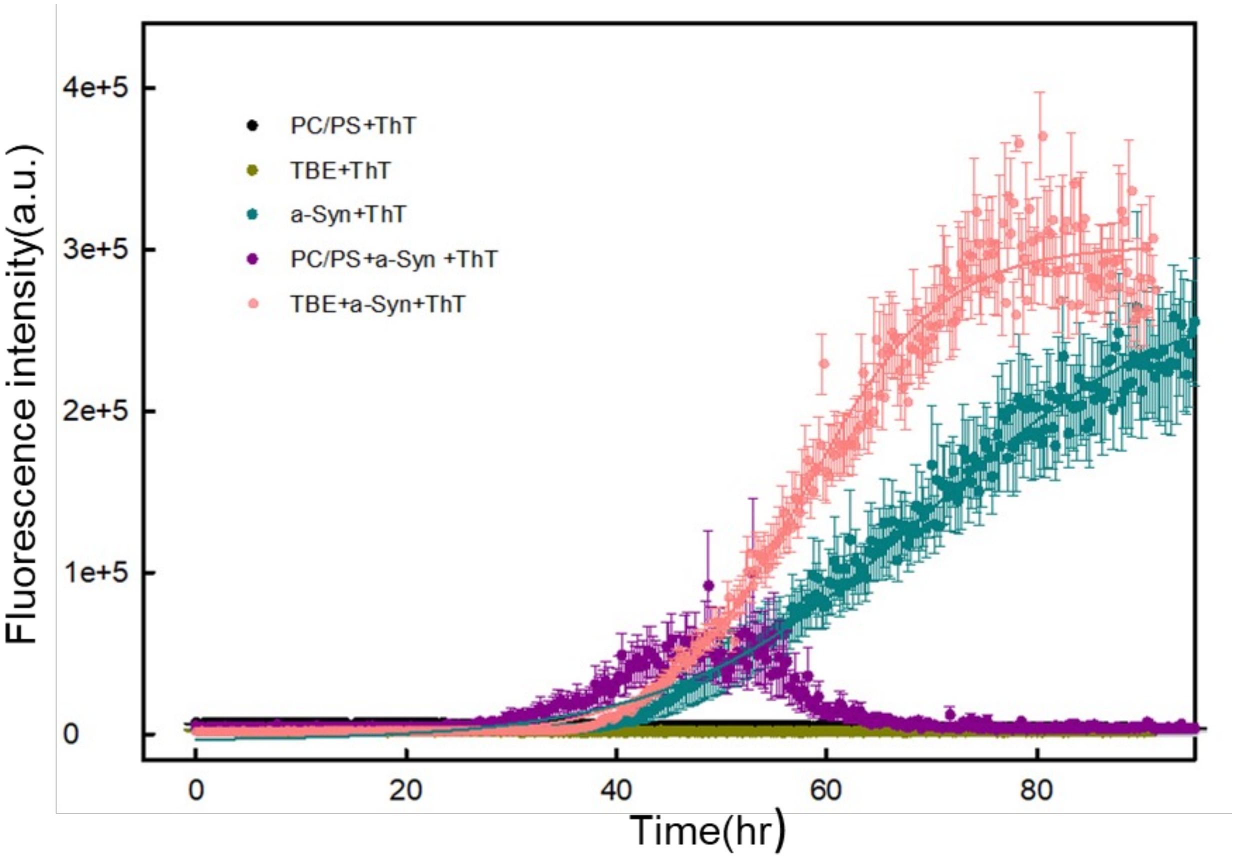


**Supplementary Figure 4: ThT fluorescence study of α-Synuclein (α-Syn) aggregation in the presence of PC/PS and TBE membrane surface.** No change was observed in the ThT fluorescence in the presence of PC:PS (6:4) and total brain extract (TBE) membrane alone (control). Significant increase in ThT fluorescence observed for α-Syn under both membrane conditions {PC:PS (6:4)} and TBE . α-Syn concentration was 50 μM. Data points were shown as the means ± S.D. of three independent measurements. All the experiments were carried out on Sodium phosphate buffer, pH 7 at 37 °C.

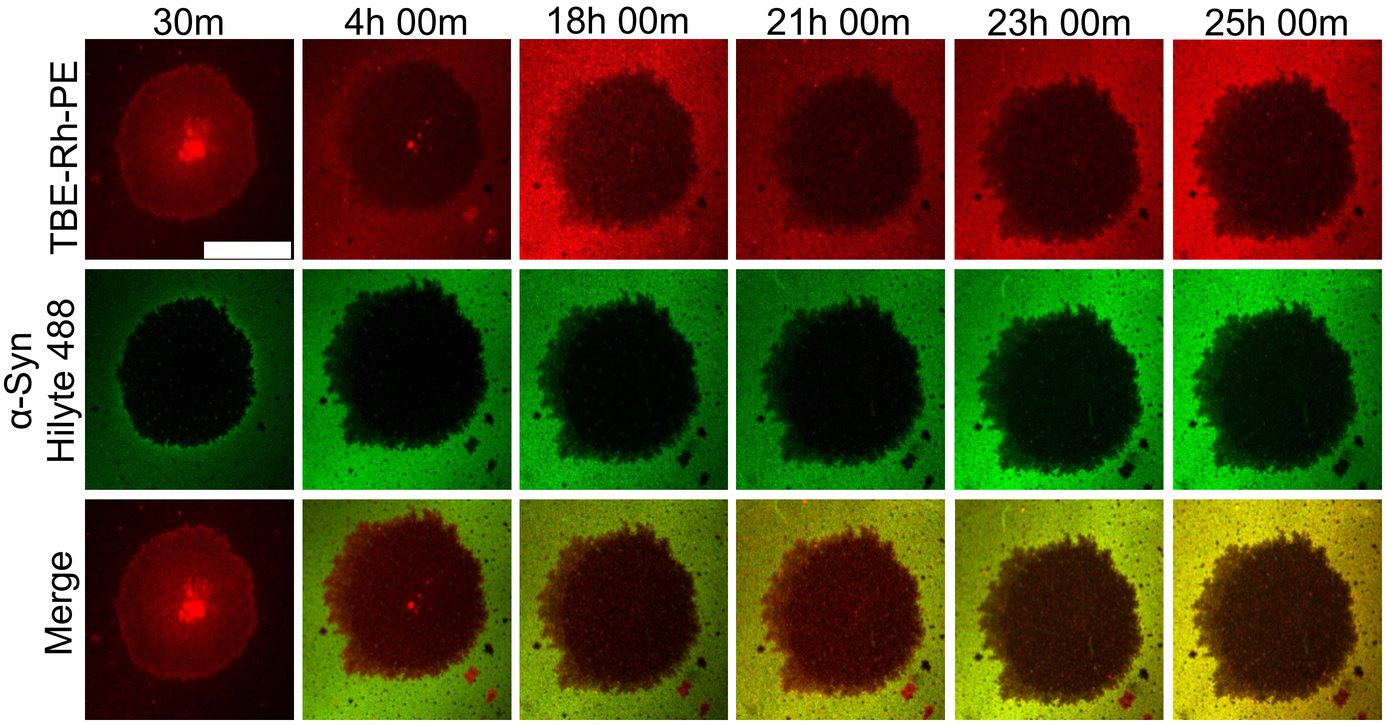


**Supplementary Figure 5: Peripheral binding of α-Syn was observed on membrane surfaces of Total brain extract (TBE).** No strong binding was observed in the presence of 100 nM α-Syn labeled with Hilyte-488 for the TBE-supported lipid bilayers (SLBs) labeled with 0.1% Rhodamine-PE (Rhod-PE) over a 25-hour period. (Scale bar, 5 μm). n = 3 independent experiments**.**

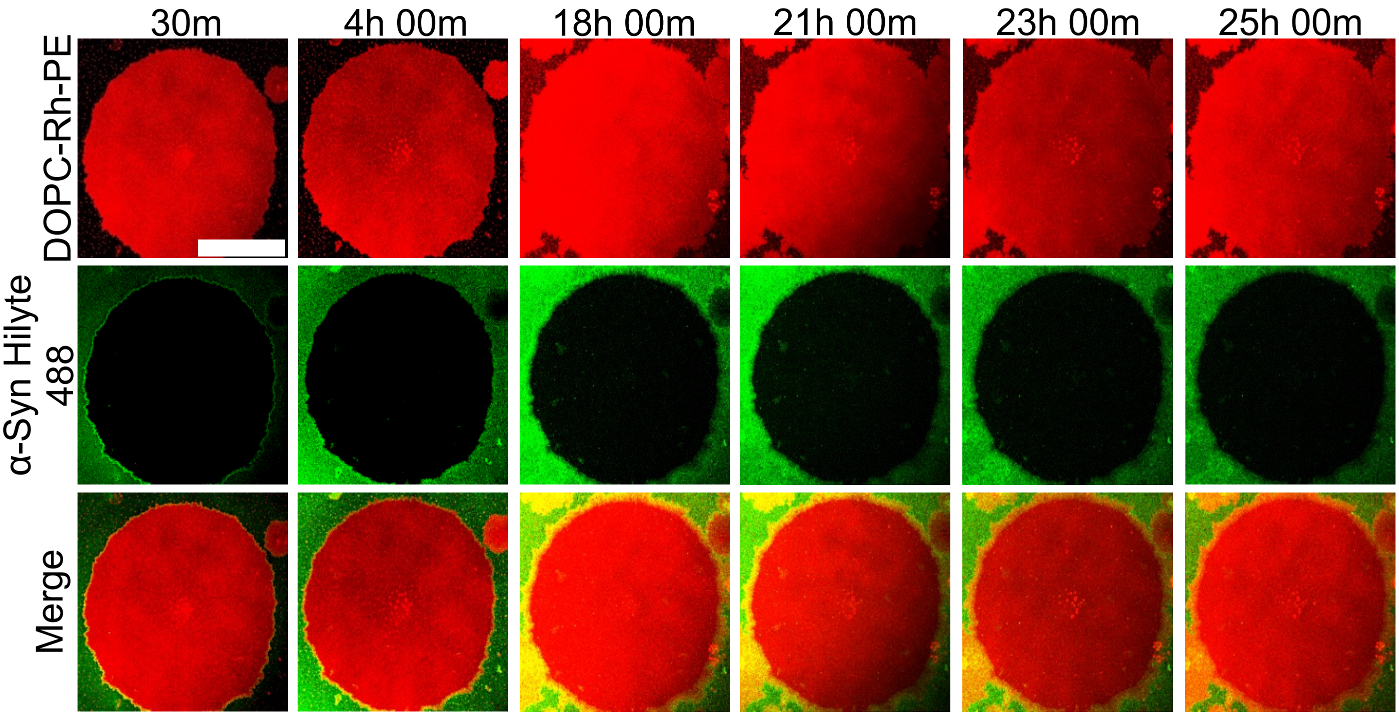


**Supplementary Figure 6: No binding of α-Syn was observed on DOPC membrane.** No binding was observed in the presence of 100 nM α-Syn labeled with Hilyte-488 for the Zwitterionic DOPC-supported lipid bilayers (SLBs) labeled with 0.1% Rhodamine-PE (Rhod-PE), over a 25-hour period. (Scale bar, 15 μM). n = 3 independent experiments.

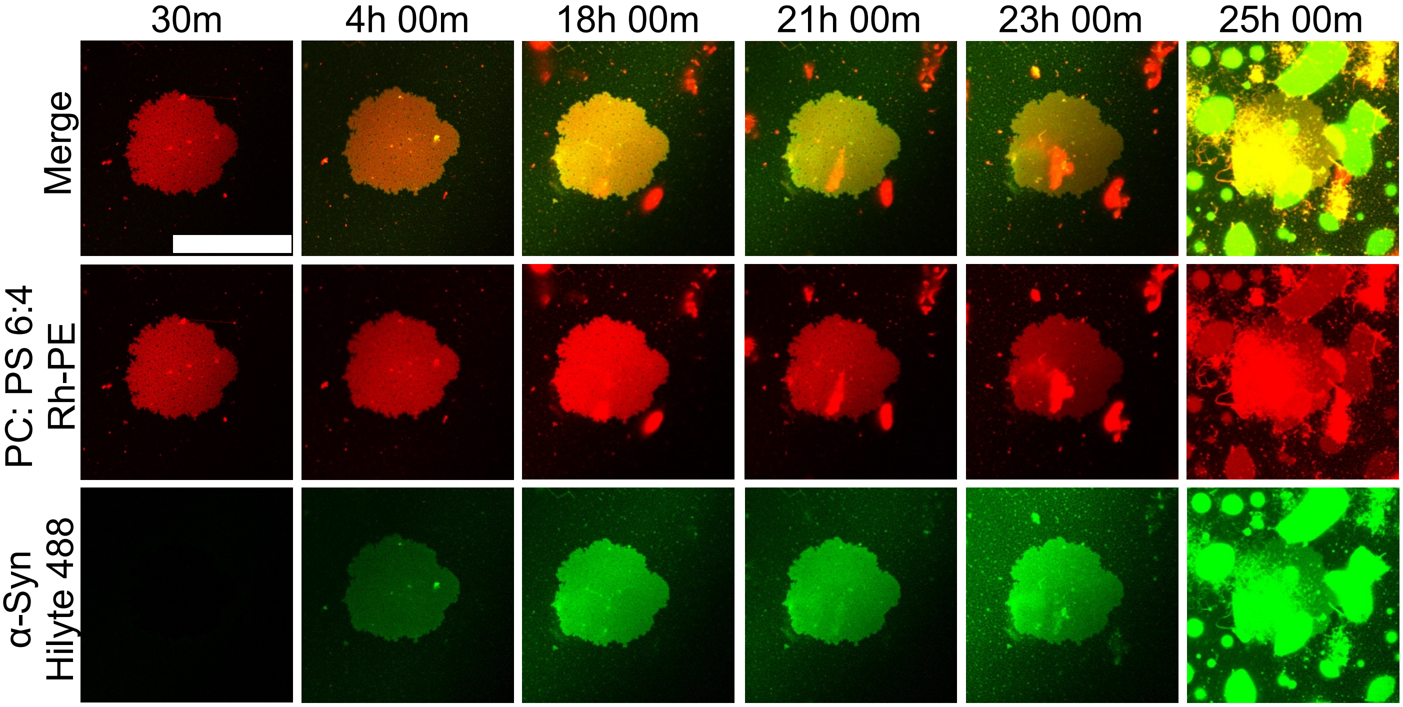


**Supplementary Figure 7: Time-Lapse Confocal Imaging of α-Syn Condensation on SLB at higher protein concentration.** Confocal microscopy imaging revealed the formation of liquid condensate in the presence of 2 μM α-Syn (Hilyte-488 labeled) on PC:PS (6:4)-supported lipid bilayers (SLBs) labeled with 0.1% Rhodamine-PE (Rhod-PE), over a 25-hour period. (Scale bar, 10 μm). n = 4 independent experiments.

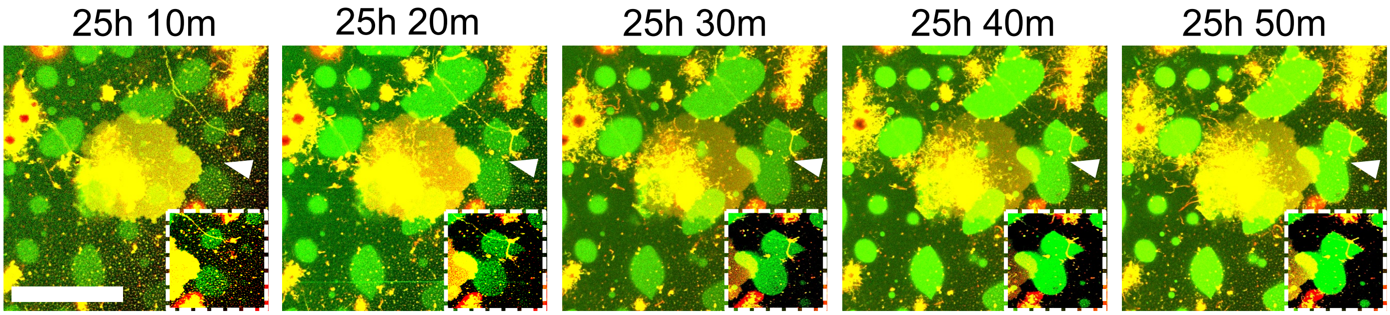


**Supplementary Figure 8: Fusion event captured in the 2 μM α-Syn concentration.** The inset and arrow highlight a fusion event that happened after 25 hours on the membrane-supported lipid bilayers composed of PC:PS (6:4) in the presence of 2 μM α-Syn. (Scale bar, 10 μm). n = 4 independent experiments.

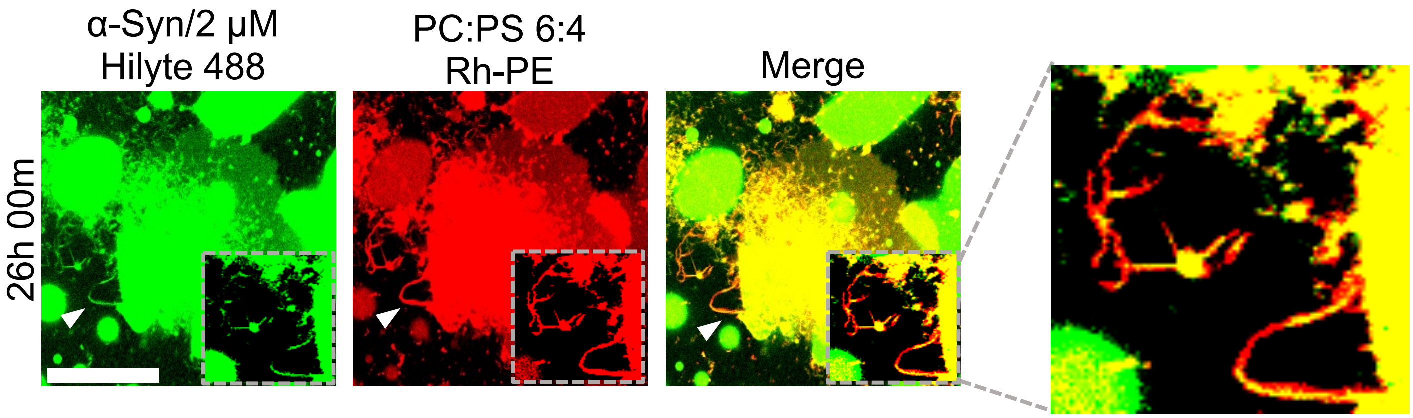


**Supplementary Figure 9: α-Syn condensation triggers fibril formation and tubulation of the membrane.** The inset and arrowhead highlight the condensate-associated fibril form after 26 hours of incubation. 2 µM α-Syn was used in the experiment. (Scale bar, 10 µm).

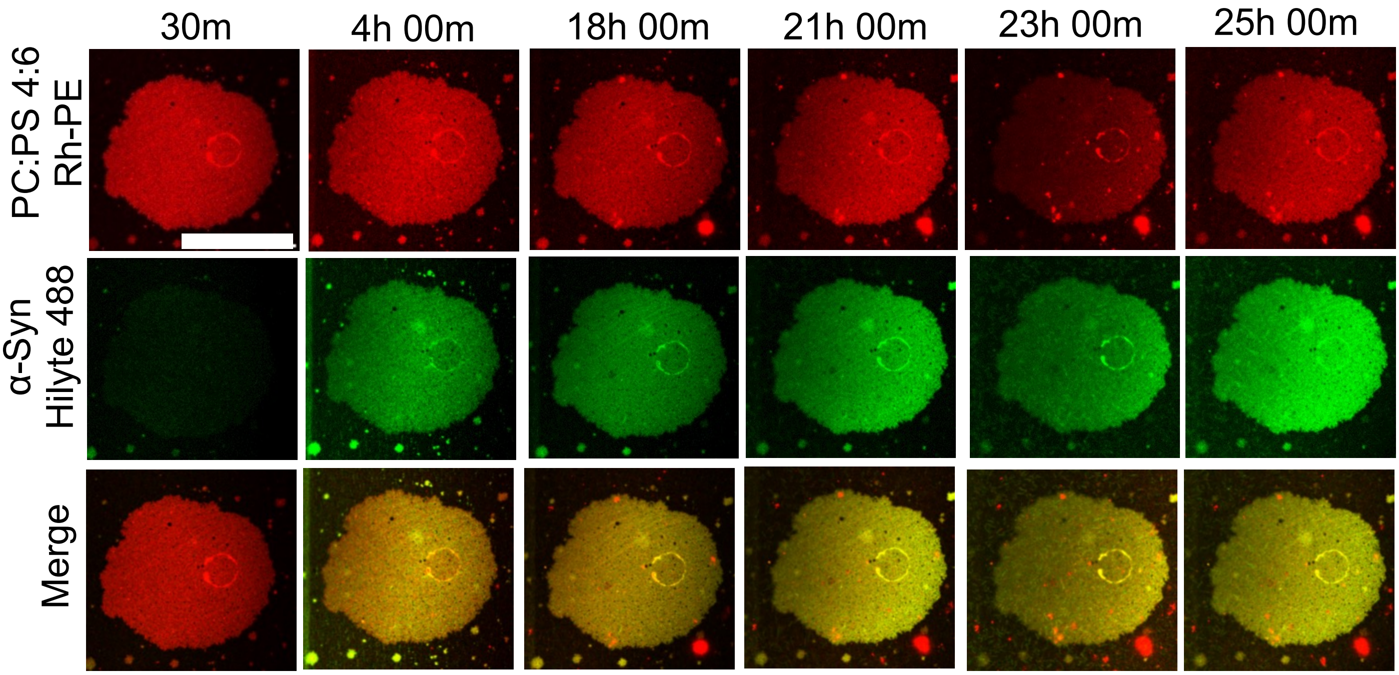


**Supplementary Figure 10: Temporal snapshots of the binding of α-Syn on PC:PS (4:6) membrane.** There was strong binding observed, but no LLPS in the presence of 100 nM α-Syn labeled with Hilyte-488 for the PC:PS (4:6)-supported lipid bilayers (SLBs) labeled with 0.1% Rhodamine-PE (Rhod-PE), over a 25-hour period. Although micellar aggregates were observed after 23 hours of incubation. (Scale bar, 10 μm). n = 3 independent experiments.

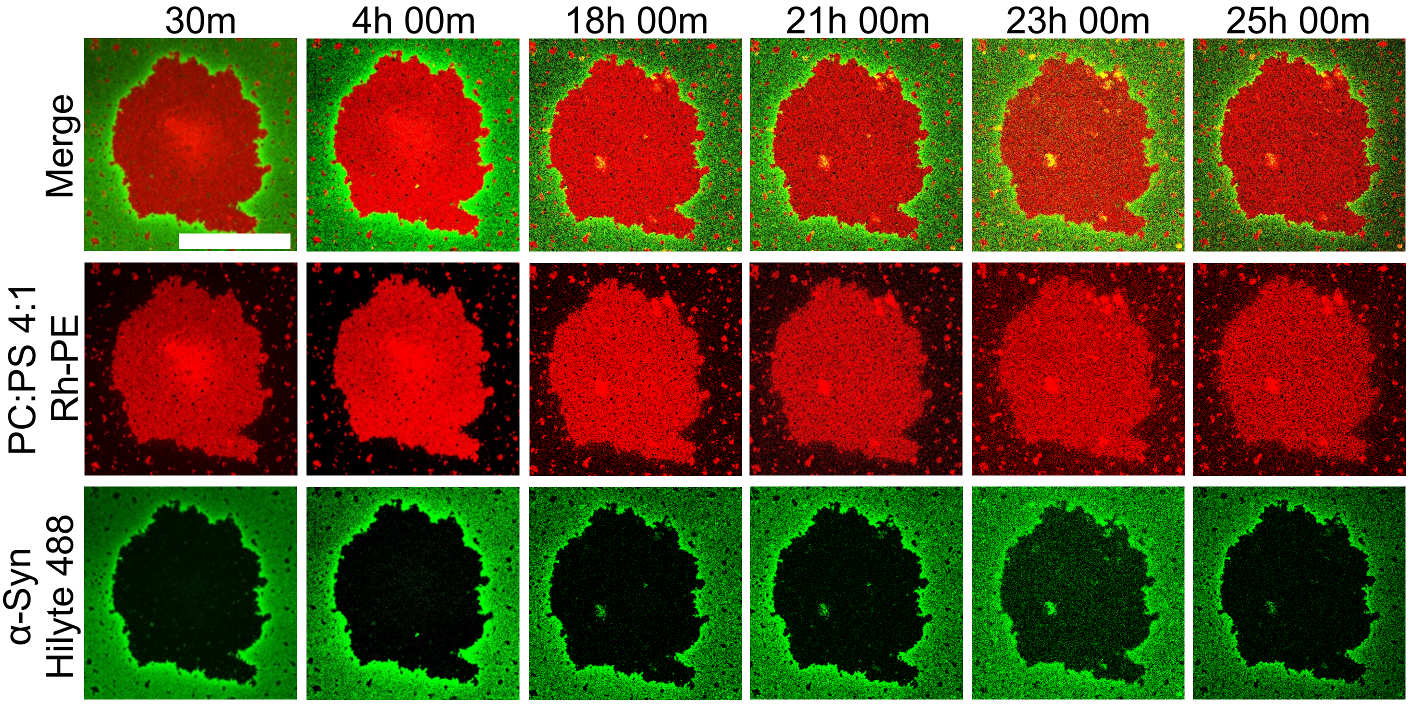


**Supplementary Figure 11: Temporal snapshots of the binding of α-Syn on PC:PS (4:1) membrane.** No binding was observed in the presence of 100 nM α-Syn labeled with Hilyte-488 for the PC:PS (4:1)-supported lipid bilayers (SLBs) labeled with 0.1% Rhodamine-PE (Rhod-PE), over a 25-hour period. (Scale bar, 10 µm). n = 3 independent experiments**.**

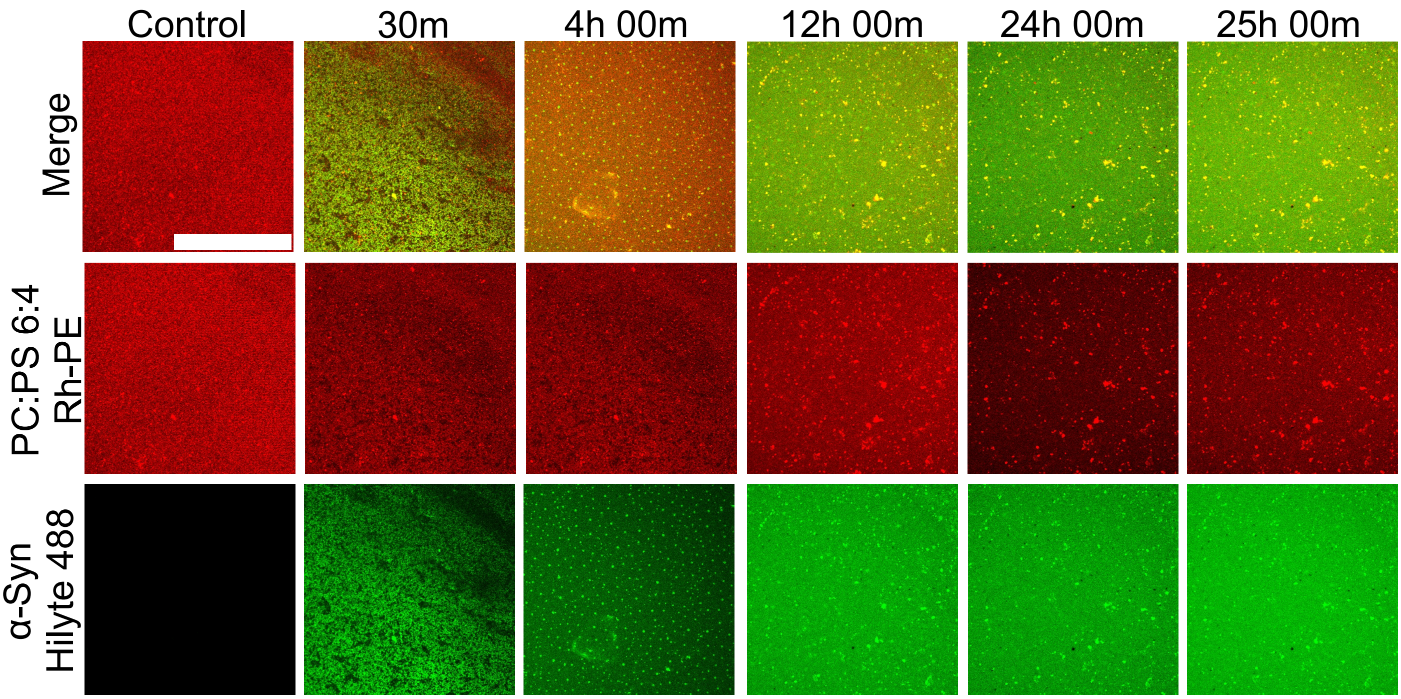


**Supplementary Figure 12: No condensation of α-Syn was observed in free lipids of PC:PS (6:4).** Confocal microscopy imaging revealed no condensate formation, but there were small aggregates observed in the presence of 100 nM α-Syn (Hilyte-488 labeled) on PC:PS (6:4)-supported lipid bilayers (SLBs) labeled with 0.1% Rhodamine-PE (Rhod-PE) over a 25-hour period. (Scale bar- 10 μM). n = 3 independent experiments.

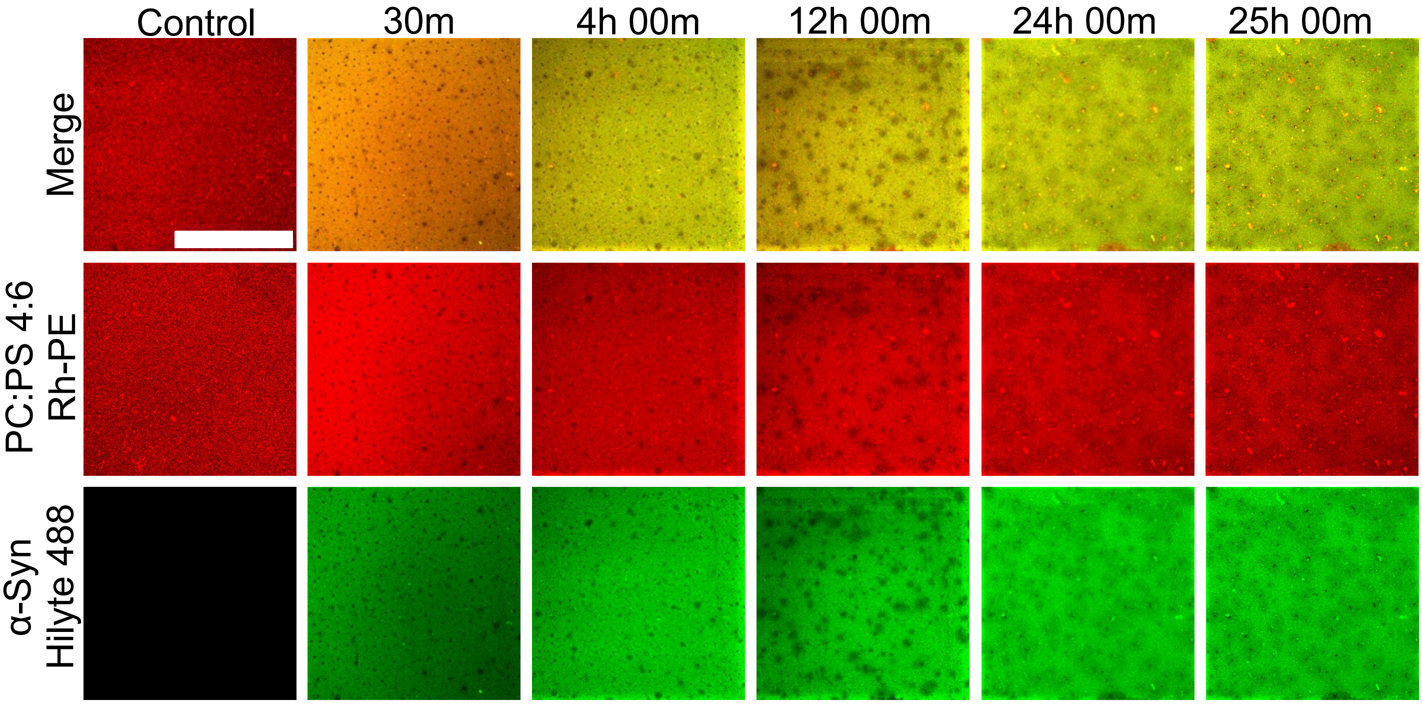


**Supplementary Figure 13: No condensation of α-Syn was observed in free lipids of PC:PS (4:6).** Confocal microscopy imaging revealed the formation of no condensate in the presence of 100 nM α-Syn (Hilyte-488 labeled) on PC:PS (4:6)-free lipids labeled with 0.1% Rhodamine-PE (Rhod-PE) over a 25-hour period. (Scale bar, 10 μm). n = 3 independent experiments.

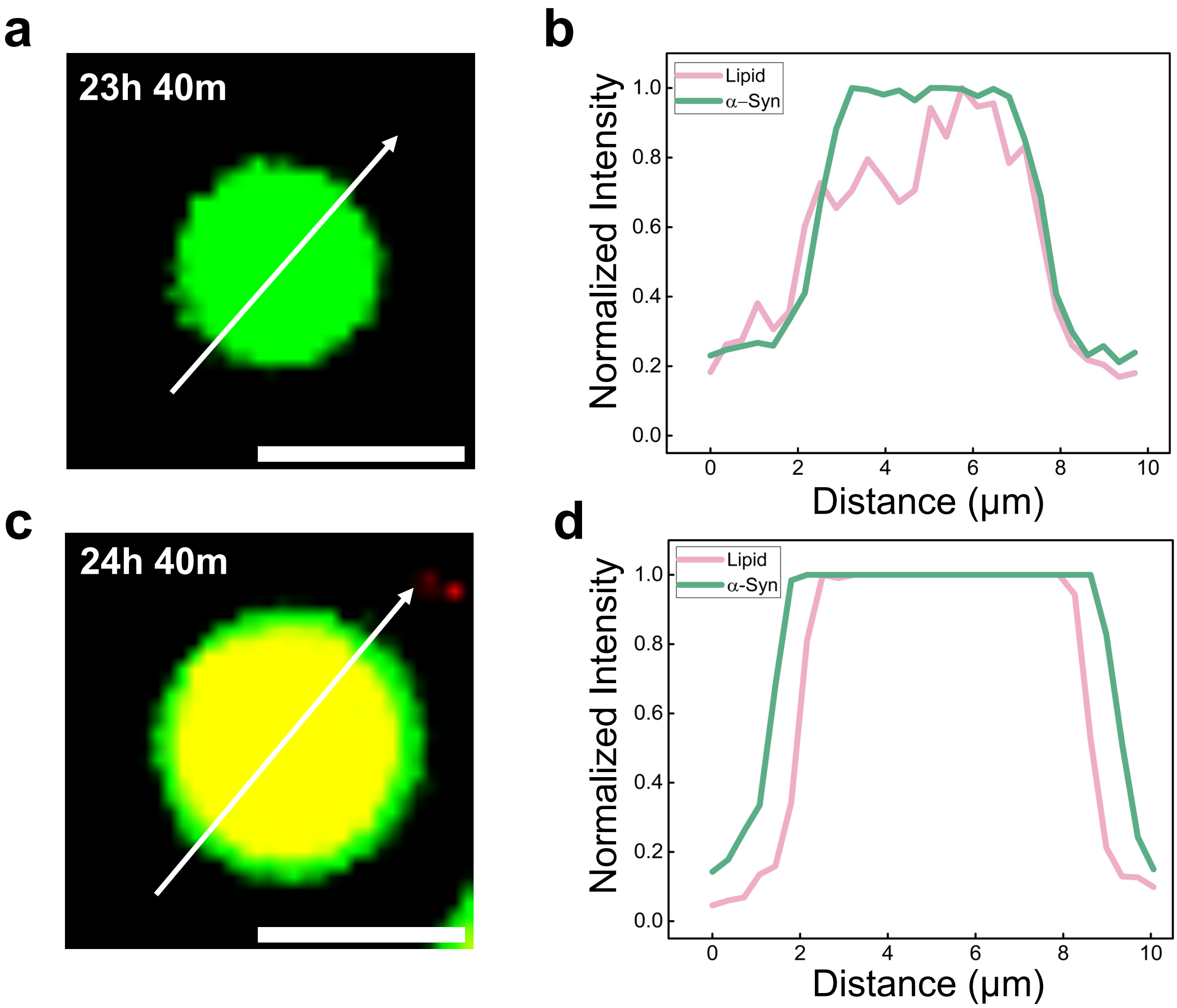


**Supplementary Figure 14: Colocalization of α-Syn and lipids within condensates.** (a and b) Colocalization of α-Syn and lipids within condensates formed at 23.40 hours and 24.40 hours (c and d) was demonstrated by confocal microscopy. 100 nM concentration of α-Syn, doped with 10% labeled with Hilyte-488, was combined with a PC:PS (6:4) membrane containing Rhodamine-PE (Rhod-PE) to visualize α-Syn and lipids, respectively. (Scale bar, 5 μm.)

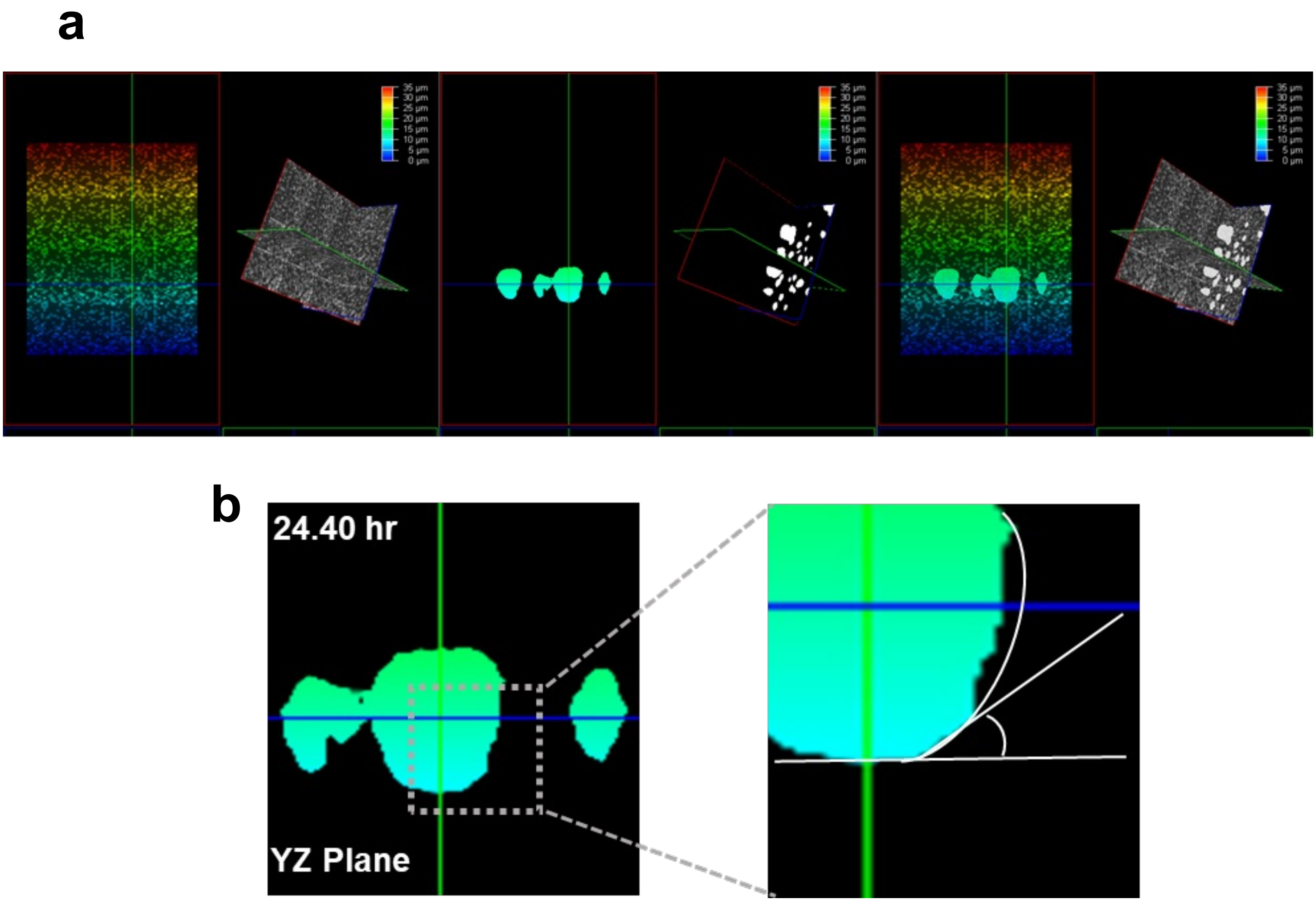


**Supplementary Figure 15: Three-dimensional analyses of α-Syn condensates on the membrane.** YZ Plane projection from a 3D image of droplets. The inlet shows the contact angle measurement of an α-Syn condensate on PC:PS (6:4) SLBs.

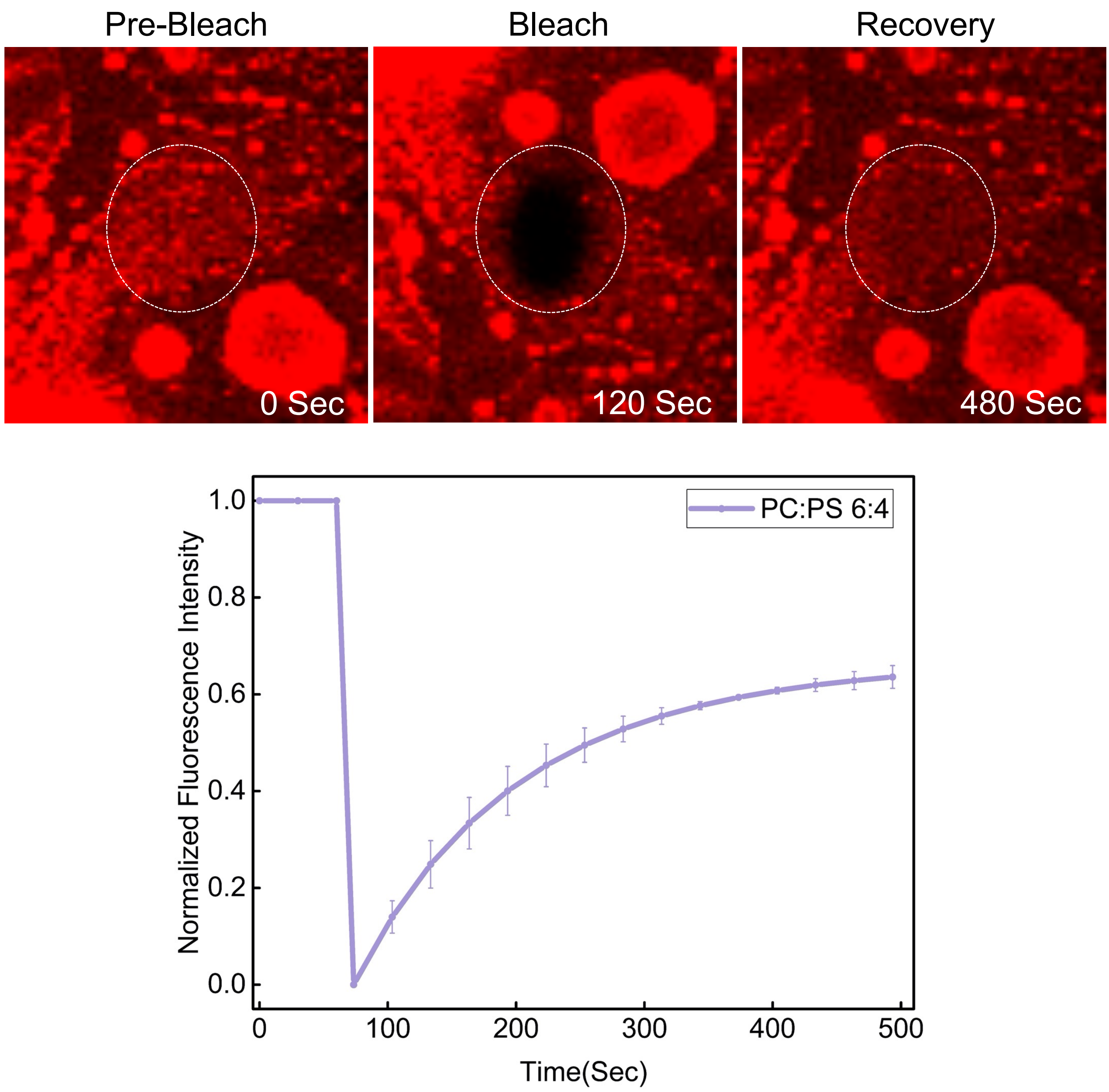


**Supplementary Figure 16: FRAP analysis of control supported lipid bilayers (SLBs) composed of PC:PS (6:4) membrane composition.** The fluorescence recovery of the membrane shows membrane fluidity on the supported lipid bilayers (SLBs) of PC:PS (6:4). n = 3 independent experiments.

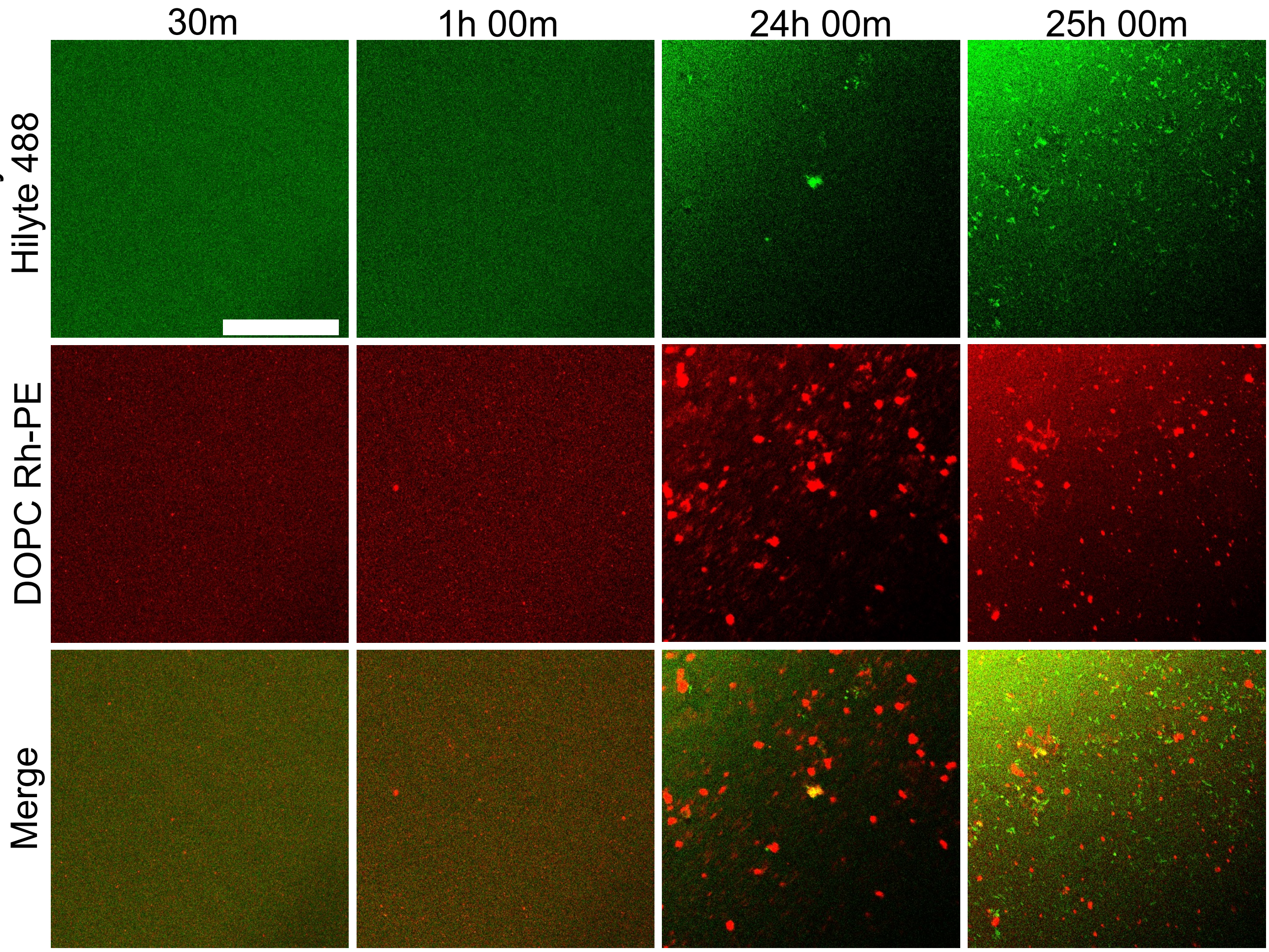


**Supplementary Figure 17: No condensation of α-Syn was observed in the presence of free DOPC lipids.** Confocal microscopy imaging revealed no condensate formation, but there were small aggregates observed in the presence of 100 nM α-Syn (Hilyte-488 labeled) in the presence of DOPC free lipids (500 μM) labeled with 0.1% Rhodamine-PE (Rhod-PE) over a 25-hour period. (Scale bar, 10 μm). n = 3 independent experiments.

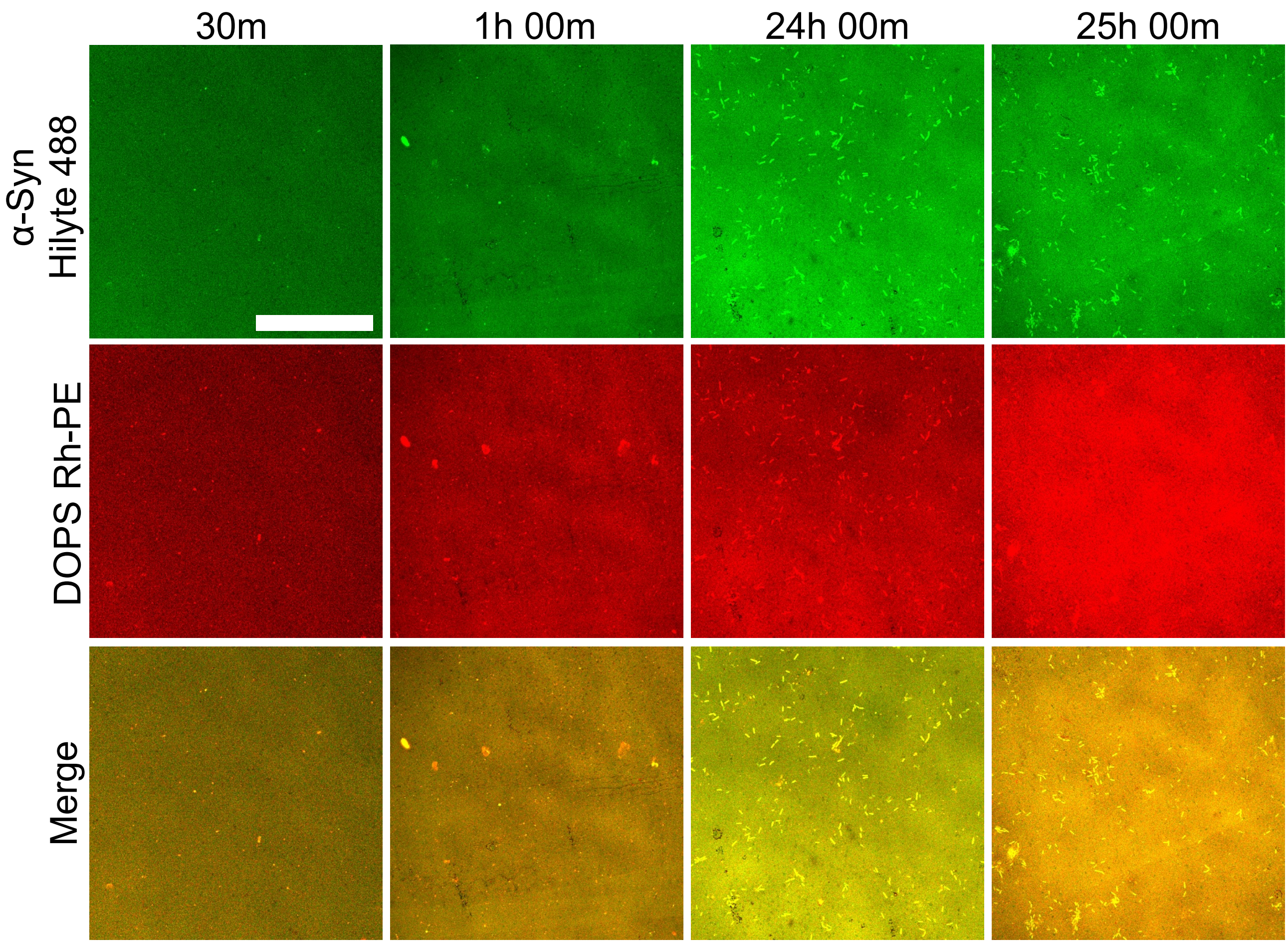


**Supplementary Figure 18: α-Syn formed fibrillar structures in the presence of free lipids of DOPS but no condensation.** Confocal microscopy imaging revealed no condensate formation, but there was a small fibrillar structure observed in the presence of 100 nM α-Syn (Hilyte-488 labeled) in the presence of DOPS free lipids (500 μM) with 0.1% Rhodamine-PE (Rhod-PE) over a 25-hour period. (Scale bar, 10 μm). n = 3 independent experiments**.**

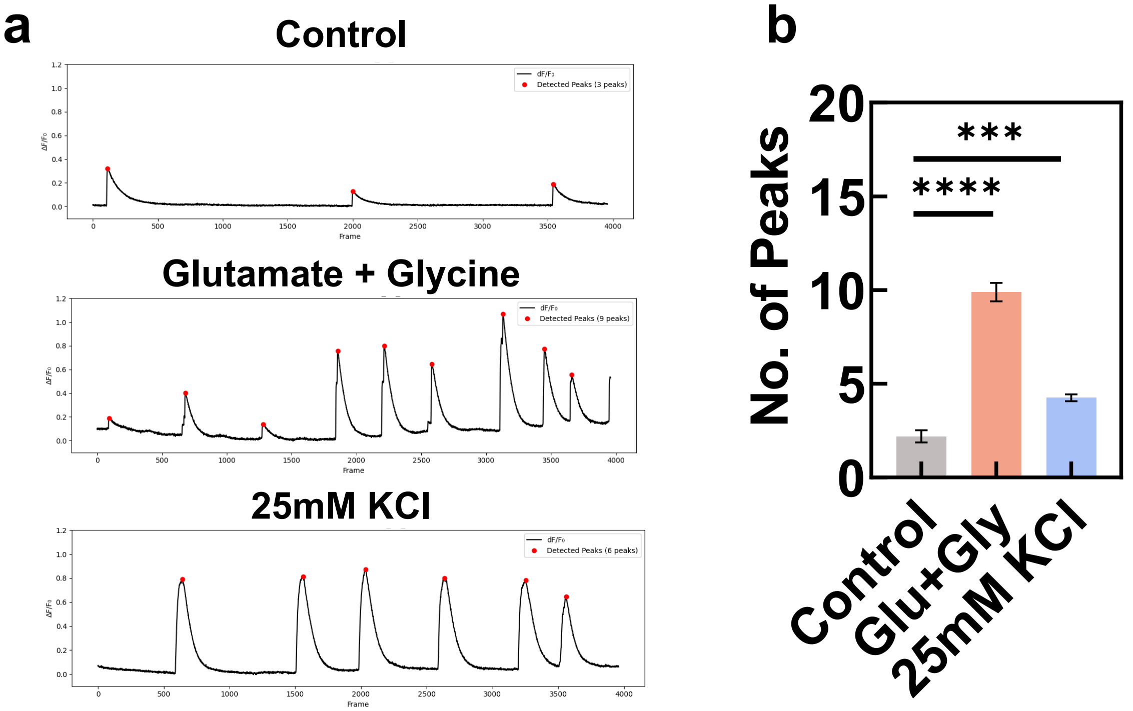


**Supplementary Figure 19: Calcium imaging confirms neuronal depolarization induced by glutamate and KCl treatment.** (a) Representative calcium imaging traces showing increased intracellular calcium levels following treatment with glutamate and KCl, indicating effective depolarization. (b) Quantitative analysis of calcium responses presented as mean ± SEM. One-way ANOVA followed by Sidak's multiple comparison test was performed between selected groups (*$p < 0.05$; **$p < 0.01$; ***$p < 0.001$; ****$p < 0.0001$).

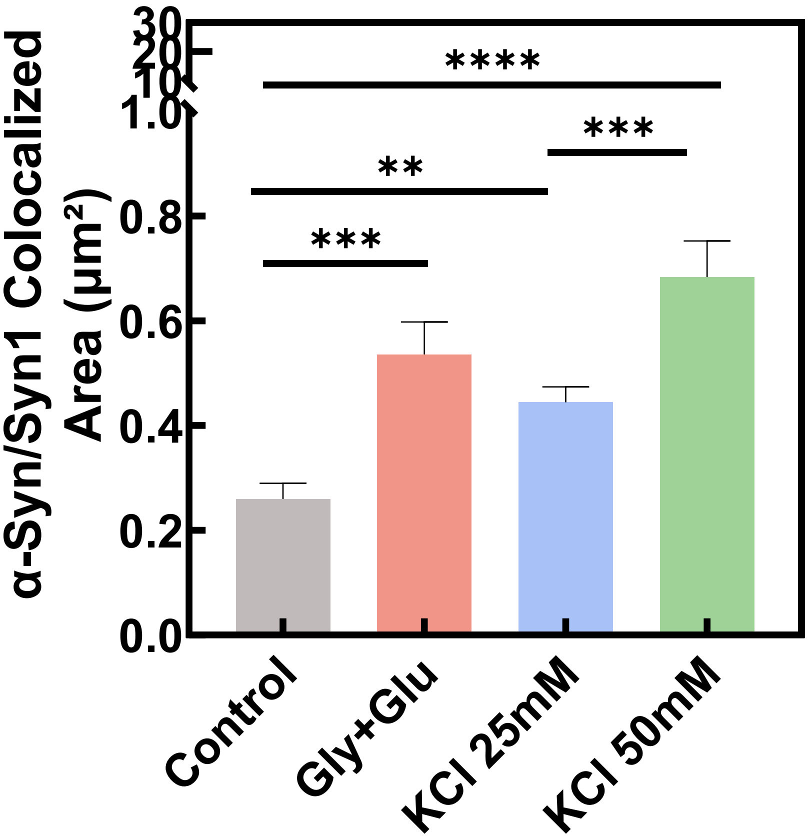


**Supplementary Figure 20: KCl stimulation increases colocalization of α-Syn puncta with the presynaptic marker synapsin-1 in a concentration-dependent manner.** Quantification demonstrates a progressive increase in the degree of colocalization between α-Syn and synapsin-1 puncta with increasing concentrations of KCl (Mean ± SD, $^{**}p < 0.01$, $^{***}p < 0.001$; $^{****}p < 0.0001$; One-way ANOVA followed by Sidak's multiple comparison test was performed between selected groups).
