## Supplementary figures and images for "Membrane Interfacial Potential Governs Surface Condensation and Fibrillation of α-Synuclein in Neurons"

### Supplementary video 2

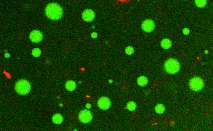
